# A Second Role for the Second Messenger Cyclic-di-GMP in *E. coli*: Arresting Cell Growth by Altering Metabolic Flow

**DOI:** 10.1101/2022.09.14.507988

**Authors:** YuneSahng Hwang, Rasika M. Harshey

## Abstract

C-di-GMP primarily controls motile to sessile transitions in bacteria. Diguanylate cyclases (DGCs) catalyze the synthesis of c-di-GMP from two GTP molecules. Typically, bacteria encode multiple DGCs that are activated by specific environmental signals. Their catalytic activity is modulated by c-di-GMP binding to autoinhibitory sites (I-sites). YfiN is a conserved inner membrane DGC that lacks these sites. Instead, YfiN activity is directly repressed by periplasmic YfiR, which is inactivated by redox stress. In *E. coli*, an additional envelope stress causes YfiN to relocate to the mid-cell to inhibit cell division by interacting with the division machinery. Here, we report a third activity for YfiN in *E. coli*, where cell growth is inhibited without YfiN relocating to the division site. This action of YfiN is only observed when the bacteria are cultured on gluconeogenic carbon sources, and is dependent on absence of the autoinhibitory sites. Restoration of I-site function relieves the growth-arrest phenotype, and disabling this function in a heterologous DGC causes acquisition of this phenotype. Arrested cells are tolerant to a wide range of antibiotics. We show that the likely cause of growth arrest is depletion of cellular GTP from run-away synthesis of c-di-GMP, explaining the dependence of growth arrest on gluconeogenic carbon sources that exhaust more GTP during production of glucose. This is the first report of c-di-GMP-mediated growth arrest by altering metabolic flow.

**Significance:** The c-di-GMP signaling network in bacteria not only controls a variety of cellular processes such as motility, biofilms, cell development and virulence, but does so by a dizzying array of mechanisms. The DGC YfiN singularly represents the versatility of this network in that it not only inhibits motility and promotes biofilms, but also arrests growth in *E. coli* by relocating to the mid-cell and blocking cell division. The work described here reveals that YfiN arrests growth by yet another mechanism in *E. coli* – changing metabolic flow. This function of YfiN, or of DGCs without autoinhibitory I-sites, may contribute to antibiotic tolerant persisters in relevant niches such as the gut where gluconeogenic sugars are found.

## Introduction

Bacteria show a remarkable ability to adapt to different environmental conditions through multiple signaling networks [1, 2]. Some of these networks commonly use cyclic or linear nucleotides to link environmental stimuli to specific bacterial responses [3, 4]. Cyclic di-GMP (c-di-GMP) was first identified in *Gluconacetobacter xylinus* for its role in cellulose production [5]. In most bacteria c-di-GMP controls the transition between motile and sessile behaviors, but is not limited to these functions [6, 7]. Broader roles for this system include virulence, progression through the cell cycle, phage resistance, and surface attachment [8–11].

Intracellular c-di-GMP levels are regulated by multiple diguanylate cyclases (DGCs) identifiable by a signature GGDEF active-site (referred to as GGDEF domain) that catalyze the synthesis of c-di-GMP from two GTP molecules, and multiple phosphodiesterases (PDEs) identifiable by EAL or HD-GYP domains that break down c-di-GMP first to linear pGpG and subsequently to GMP [12, 13]. Although ectopic expression of most DGCs and PDEs often leads to dramatic phenotypes, their native expression often does not, as observed by lack of an observable phenotype when their genes are experimentally mutated, indicating that these enzymes may be only activated under specific environmental conditions. The presence of specific sensory modules such as HAMP, PAS, LuxR or BLUF domains allows these enzymes to respond to a variety of sensory cues, both chemical and physical; however, most signals activating these enzymes are still unknown [14–18]. The c-di-GMP produced in response to these signals elicits its varied outputs primarily by binding to cellular effectors and targets.

DGCs are regulated not only by environmental signals but can also self-regulate. An autoinhibition (I) site located close to the active site was first identified in the DGC PleD of *Caulobacter crescentus*, where it regulates enzyme activity through negative feed-back, curtailing excess c-di-GMP production [12]. This site (primary I site; I_p_) is identifiable by an RXXD consensus located five amino acids upstream of the GGDEF motif. c-di-GMP bound at I_p_ causes a conformation change that leads to the repression of enzymatic activity; accordingly, mutations in conserved I_p_ residues increase c-di-GMP production [12]. Based on structural studies, additional arginine residues located outside the GGDEF domain in PleD were also seen to contribute to the c-di-GMP pocket for feedback inhibition; these were designated the secondary I site (I_s_) [19]. A different role for I_s_ was reported for the DGC GcbC in *Pseudomonas fluorescens*, where this site is required for interaction of GcbC with its target protein LapD, which controls biofilm production [20].

YfiN, also called DgcN or TpbB, is a conserved inner membrane DGC with a Per-Arnt-Sim (PAS)-like sensory domain in the periplasm and HAMP-GGDEF domains in the cytoplasm [21, 22]. *E. coli* YfiN lacks both I_p_ and I_s_ sites, as do the majority of bacterial YfiN homologs. In *E. coli, Salmonella enterica* and *Pseudomonas aeruginosa*, the enzymatic activity of YfiN is repressed by periplasmic YfiR [16, 23, 24]. Redox stress can cause YfiR to dissociate from YfiN, activating its DGC function [16, 24]. In *P. aeruginosa*, transposon disruption of *yfiR* led to increased biofilm formation through the Pel/Psl biosynthesis pathway, and enzymatically inactive YfiN showed less virulence in the mouse model compared to wild-type (WT) [16]. YfiN was also shown to be important for biofilm maintenance in response to peroxide stress in *P. aeruginosa* [25].

In *E. coli*, YfiN exhibits the canonical motility-inhibition function of a DGC [24]. A second very different function for YfiN is a two-step response to redox and envelope stresses: in response to redox stress YfiN produces c-di-GMP, and in response to a second envelope stress YfiN moves to the mid-cell to arrest cell division [24]. Unlike known cell division inhibitors [26], the interaction of YfiN with cell division proteins FtsZ and ZipA retains the Z ring at the mid-cell but prevents septal invagination [24]. We report here a third function for YfiN in *E. coli,* where YfiN arrests growth by a mechanism different from cell division arrest. This mechanism exploits the absence of regulatory I sites in this DGC. We show that when growing on gluconeogenic sugars, unregulated c-di-GMP synthesis by YfiN depletes cellular GTP, which is the likely cause of failure of all macromolecular synthesis. Growth arrest can be reversed by reintroducing the I sites. Such a function for a DGC has not been previously reported in bacteria. This ability of YfiN to enable survival by shutting down growth, allows the bacteria to tolerate a broad spectrum of antibiotics.

## Results

### YfiN arrests *E. coli* growth on specific carbon sources

Prior studies conducted in *E. coli* growing in LB with ectopically expressed YfiN reported that even though expression was induced from the beginning of a growth cycle, YfiN appeared at the division site just prior to the onset of the stationary phase, resulting in cell division arrest [24]. When cells were grown in minimal M9 glycerol (M9M) however, we observed that irrespective of whether the ectopic expression was from a plasmid (pYfiN; PBAD promoter) or from its chromosomal location (cYfiN_GFP_; PTrc promoter; all cYfiN experiments were with the GFP-tagged version and will be referred to as cYfiN henceforth), growth began to slow as early as two hours post-YfiN induction and to remain arrested (**Fig. 1A, B**); western blot analysis showed cYfiN levels to be ∼8 fold lower than pYfiN (**Fig. S1A**). The DGC function of YfiN was important for this phenotype, as observed by lack of growth arrest in the active site mutant pYfiN(GGAAF), where the GGDEF signature motif is changed to GGAAF **(Fig. 1C**); this mutation did not compromise protein stability (**Fig. S1B**).

**Figure 1.**
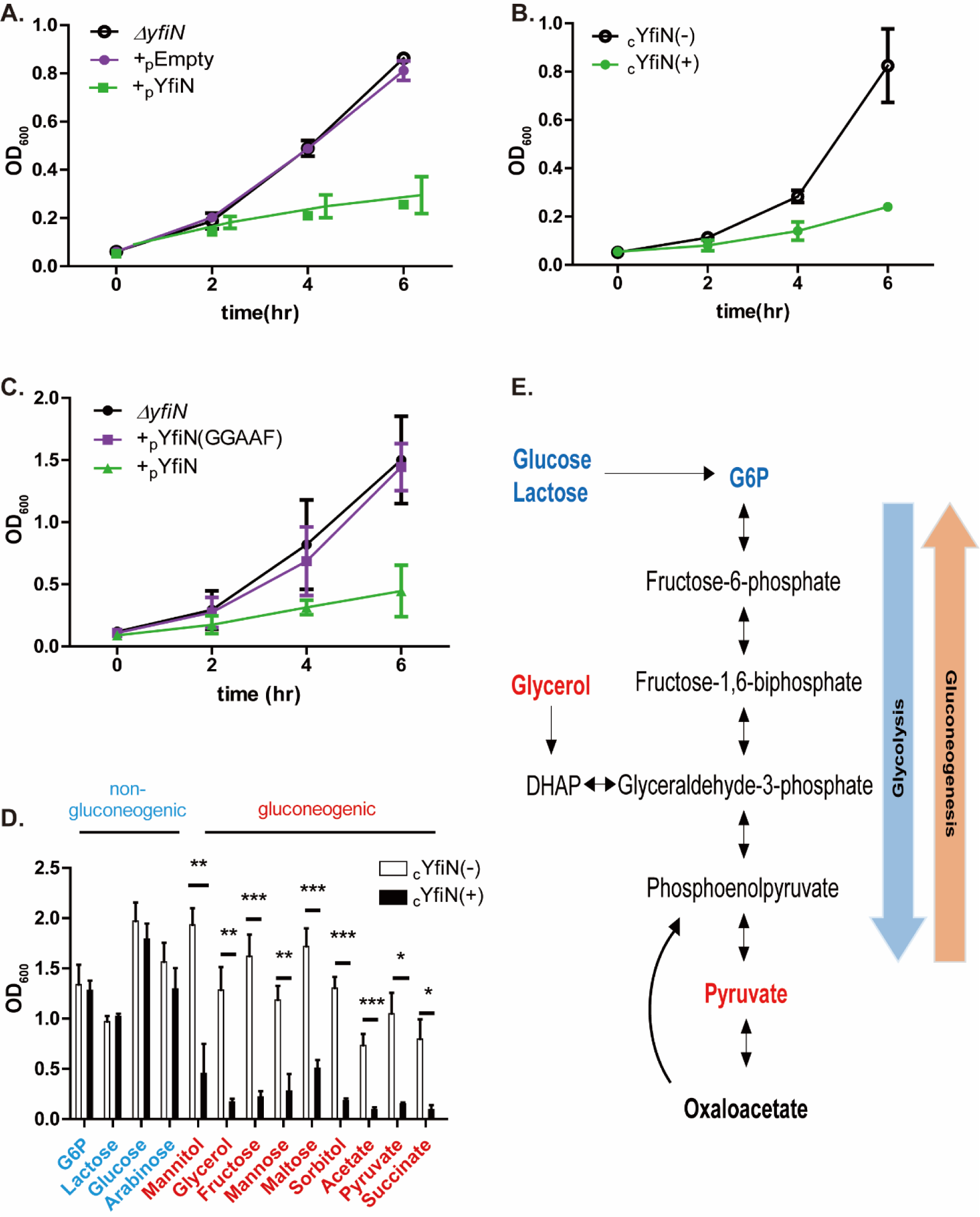
YfiN arrests *E. coli* growth only on specific carbon sources. *(A)* Time course of growth (OD600) in M9M of Δ*yfiN E. coli* harboring either pYfiN or pEmpty (control) plasmids, where expression of *yfiN* is driven from the PBAD promoter (see Methods for media and arabinose inducer concentrations). *(B)* As *(A)*, except *yfiN* expression is from its chromosomal location (cYfiNGFP), driven from the PTrc promoter. +/-indicate IPTG addition. *(C)* As in *A*, except activity of pYfiN is compared with its active site mutant YfiN(GGAAF). (*D)* As in *B* (cYfiN), except cells were grown in M9 media supplemented with indicated sugars, and OD600 measured at 8h. (*E*) Schematic of the gluconeogenesis and glycolysis pathways, with relevant metabolites color-coded as in *D*. Data in all experiments, here and in the following figures (where applicable), were analyzed with the unpaired t-test, with three biological replicates for each sample, unless noted otherwise. p-value <0.001 ***, <0.01 **, and <0.5 *.

To test whether localization of YfiN to the mid-cell is required for the growth arrest observed in Figure 1 (**A,B**), the location of both cYfiN and pYfiN_GFP_ was monitored in M9M. cYfiN was found dispersed throughout the cell, while pYfiN_GFP_ relocated to the mid-cell in 60% of the cells (**Fig. 2A**). Thus, the observed growth-arrest by cYfiN is not related to its ability to localize at the division site, while that of pYfiN_GFP_ may include division arrest (**Fig. 1A, B**). To investigate if the mid-cell localization of pYfiN_GFP_ in 60% of the cells was related to the higher expression levels from a plasmid, we monitored the behavior of a plasmid encoded R260A mutant of YfiN (R260 is located in the cytoplasmic domain) that does not localize to the mid-cell (**Fig. 2B;** see **Fig. S1B** for protein expression). Like pYfiN_GFP_, pYfiN(R260A) _GFP_ induction resulted in severe motility inhibition on soft agar plates, indicating robust c-di-GMP production (**Fig. 2C**). Despite its failure to localize to the mid-cell, the R260A mutant arrested growth similar to WT YfiN (**Fig. 2D**). We conclude that the inability to grow in M9M is a new phenotype of YfiN, distinct from its ability to arrest cell division by interacting with the divisome at the mid-cell.

**Figure 2.**
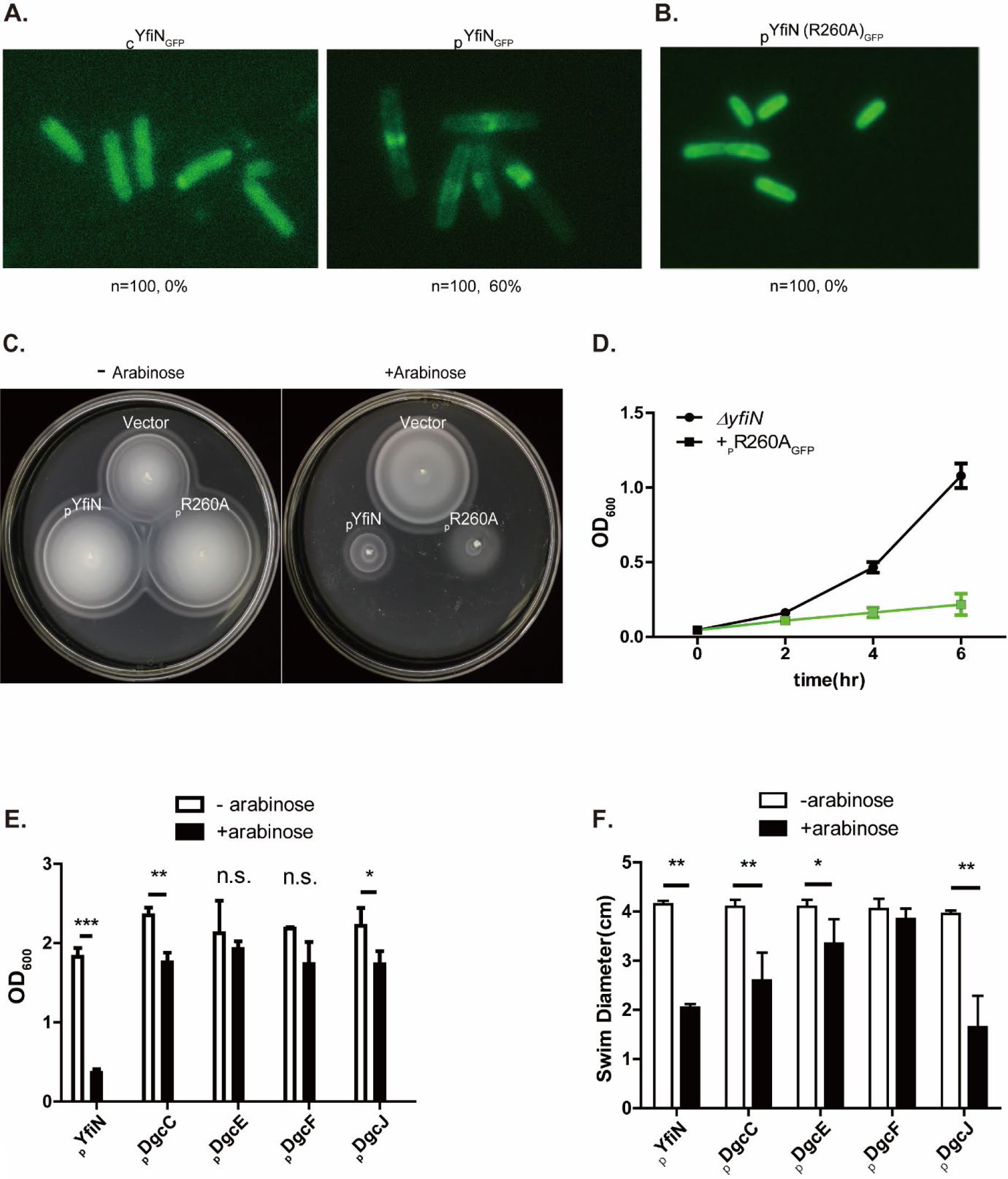
Growth-arrest phenotype of YfiN in M9M is independent of its relocation to the mid-cell. *(A)* Fluorescence images of cYfiNGFP and pYfiNGFP expressed for 4 hours in M9M. The percentage of cells with YfiNGFP foci at the mid-cell is indicated, with n = total number of cells observed. *(B)* As in *A*, except with pYfiN (R260A)GFP. *(C)* Motility of indicated strains: empty vector, pYfiNGFP, pYfiN(R260A)GFP,-/+ arabinose inducer in 0.3% soft agar plates. *(D)* As in *C*, except growth curves of the indicated strains in M9M. *(E)* A panel of all five inner membrane DGCs of *E. coli* were cloned in the PBAD vector and expressed for 12 hoursprior to recording OD600. (*F*) Motility assay of strains from *E* in soft agar plates.

*E. coli* has 12 DGCs, of which five (including YfiN) harbor a transmembrane region. To test if this new phenotype of YfiN is related to its membrane location, and to also control for overexpression artifacts, we cloned the other four transmembrane DGCs (C, E, F and J) [27] under PBAD control. In contrast to YfiN, none of these four DGCs arrested growth in M9M (**Fig. 2E**). Except for DgcF, the ectopic expression of DgcC, E and J inhibited motility, suggesting that they were active as DGCs (**Fig. 2F)**, which is supported by previous work [28–30]. We conclude that early growth arrest in M9M is unique to YfiN.

A clear difference between the early growth-arrest observed in this study (**Fig. 1A,B**) and the late growth-arrest phenotype of YfiN reported earlier [24] was the growth medium: minimal M9M in the present study, and nutrient-rich LB in the earlier one. When glycerol was substituted with glucose, the most favored carbon source for *E. coli*, cYfiN no longer inhibited growth in M9M (**Fig. 1D**, third set of bars from the left). In *E. coli* growing aerobically, glucose is directly integrated into glycolysis as glucose 6-phosphate (G6P) and consumed through the tricarboxylic acid cycle (TCA), while glycerol, an energy-poor carbon source is incorporated into central metabolism as dihydroxyacetone phosphate (DHAP), a metabolite that can participate in both gluconeogenic and glycolytic processes [31] (**Fig. 1E**). To explore this finding further, we provided several gluconeogenic and non-gluconeogenic substrates as carbon sources in the M9 medium. For the former we chose G6P, lactose, glucose and arabinose, and for the latter we chose the following: sugar alcohols in addition to glycerol (sorbitol, mannitol), various other sugars (fructose, mannose, maltose), acetate, and TCA cycle intermediates (pyruvate, succinate). There was a clear-cut difference in the ability of YfiN to arrest growth on the two sets of carbon sources, which was patterned after the glycerol and glucose examples i.e. only gluconeogenic sugars arrested growth. If the differential growth arrest phenotype of YfiN on these two types of substrates is because of the higher energy produced by one substrate type versus the other, the expectation is that providing both substrates would override the effect of the gluconeogenic substrate. This was found to be the case (**Fig. S2A-D**). To test whether YfiN-mediated growth arrest is reversible, either glucose or LB was added 2 hours post-induction of YfiN. Bacteria resumed growth immediately, suggesting that YfiN-mediated growth arrest is fully reversible and that cells are not dead (**Fig. S2E**).

### Growth arrest is correlated with absence of autoinhibitory I sites in YfiN

Given that the DGC activity of YfiN is required to mediate growth arrest (**Fig. 1C**), we first tested whether the canonical function of c-di-GMP (turning on biofilm pathways / shutting down motility) played a role. This was done in two ways: 1) simultaneous expression of YfiN with YhjH (the most active PDE in *E. coli*) expected to degrade c-di-GMP and 2) disruption of six key genes that control the major biofilm pathways in *E. coli* (production of cellulose, PGA, colanic acid, Type 1 fimbriae, EPS etc.) [32]. In the first case, i.e. expression of YhjH, the inhibitory effect of YfiN on motility was substantially relieved (**Fig. S3A**); the reduction in c-di-GMP levels was confirmed using a riboswitch-based biosensor (see Methods) [33, 34] (**Fig. S3B**). Yet, the growth arrest function of YfiN was not relieved (**Fig. S3C**). In the second case, i.e. disruption of multiple biofilm pathways, biofilm levels decreased as expected (**Fig. S3D**). However, YfiN still arrested growth (**Fig. S3E**). In summary, the canonical functions of c-di-GMP do not contribute to the observed growth arrest phenotype of YfiN.

In thinking about what distinguishes YfiN from the all the other inner membrane *E. coli* DGCs previously tested (Dgc C, E, F, J), we noted that YfiN lacks the consensus autoinhibitory I site residues (I_p_ and I_s_), which bind c-di-GMP to feedback regulate enzyme activity (**Fig. 3AB**) [12, 35]. We reasoned that lack of autoregulation would lead to continual c-di-GMP production, hence depletion of cellular GTP. Depletion of GTP as a mechanism of growth arrest is well-established in the (p)pGpp-mediated pathway [36]. Imbalance in nucleotide pools is also known to disrupt growth [37, 38]. To test the GTP depletion idea, we rebuilt the two YfiN I sites (I _p_ and I_s_) both sequentially and together. I_p_ was restored by changing GLRH to the consensus RXXD, and I_s_ by changing N280 to R. Only when both I sites were reconstituted, was growth arrest fully reversed (**Fig. 3C**). These changes did not perturb either the DGC activity of the I-site reconstituted strains as monitored by motility inhibition in soft agar plates (**Fig. 3D**) or by protein expression levels (I_ps_ mutant; **S1B**). We also measured c-di-GMP levels in YfiN and its mutants using the riboswitch-based biosensor. As expected, introduction of I site mutations reduced enzymatic function (I_p_ and I_s_), with I_ps_ showing the lowest c-di-GMP levels (**Fig. S4**). To test whether the I-site is integral to the growth arrest phenotype, we mutated the I_p_ site (RESD → GESD) in DgcA, a robust heterologous DGC from *C. crescentus* [12]. Expression of WT DgcA in *E. coli* did not elicit growth arrest in M9M, but expression of the DgcA I_p_ mutant did (**Fig. 3E**). We note that c-di-GMP levels in DgcA were similar to those when YhjH was co-expressed with YfiN (**Fig. S3B**), yet growth arrest was seen in the latter i.e. YfiN + YhJH (**Fig. S3C**) but not the former i.e. DgcA (**Fig. 3E**). Thus, both loss of the growth-arrest phenotype in YfiN and gain of this phenotype by DgcA by merely restoring and inactivating I-site function, respectively, allows us to conclude that unregulated c-di-GMP production in the absence of the I site is responsible for the arrest not by increasing c-di-GMP levels per se, but likely by depleting cellular GTP levels through continuous c-di-GMP synthesis.

**Figure 3.**
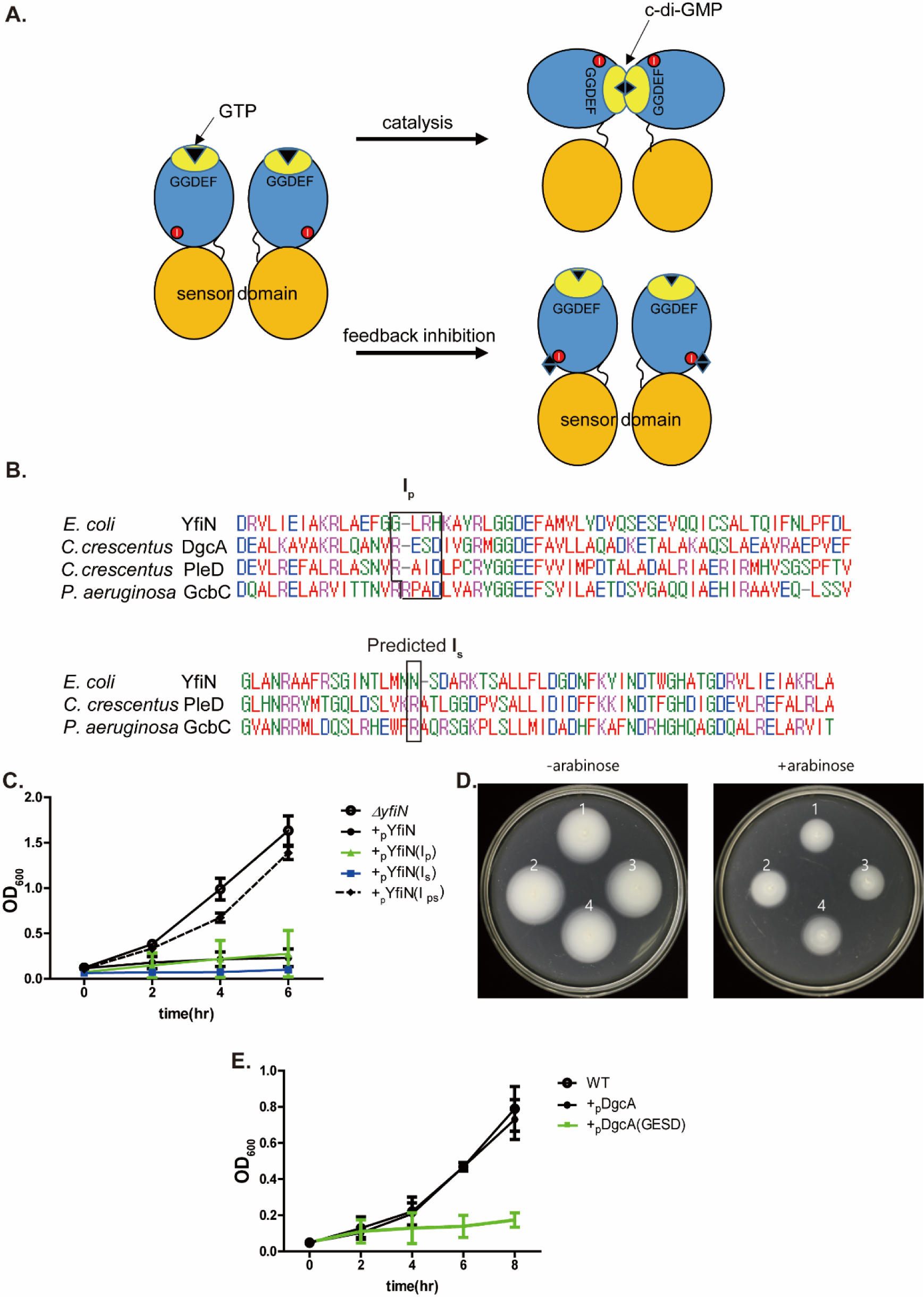
Growth arrest by YfiN is related to absence of autoinhibitory I sites. *(A)* Schematic of I-site mediated feedback inhibition. Dimeric DGCs sense environmental stimuli though their sensor domains (orange), and relay the signal to their catalytic domain (blue) to synthesize c-di-GMP. Upon reception of the sensory signal, GGDEF active sites (yellow) align in the dimer [7] to produce c-di-GMP (diamond) from 2 GTPs. Product inhibition occurs when c-di-GMP binds to I-sites (red circles), preventing the two active sites from interacting. (*B*) The primary (Ip) (RxxD) and predicted secondary (Is) auto-inhibitory sites are boxed in a sequence alignment of various DGCs around these sites (see Methods). (*C*) Growth in M9M of pYfiN derivatives reconstituted for primary (Ip), secondary (Is) or both (Ips) I-sites. Expression was induced with arabinose. (*D*) Motility of indicated strains (1: pYfiN, 2: pYfiN Ip, 3: pYfiN Is, 4: pYfiN Ips) assayed in soft agar plates. (*E)* As in C, except with cells expressing DgcA and its mutant Ip site (RESD◊GESD).

### YfiN expression depletes intracellular GTP

To test our conjecture that GTP depletion due to unregulated c-di-GMP synthesis is the cause of growth arrest by YfiN, we first compared cellular nucleotide levels with (+) or without (-) induction of _c_YfiN by metabolomic analysis of a standard panel of 219 intracellular metabolites. Of these, the levels of five ribonucleotides relevant to this study are shown in **Figure 4AB**. c-di-GMP levels were high in the cYfiN+ strain as expected. A consistent decrease in the levels of GTP, ATP, UTP and CTP was seen in experimental samples compared to the controls. To confirm the drop in GTP concentration upon cYfiN expression, we next used a luciferase-linked GTP assay (GTPase-Glo™), which converts GTP into ATP, the latter detected by the light produced by the luciferase reaction (**Fig. 4C**). We observed a 50% decrease in GTP levels in cYfiN-expressing cells, supporting the GTP metabolomics data. To monitor GTP levels by a third method, as well as to test the proficiency of various YfiN I-site mutants in converting GTP to c-di-GMP, cell lysates expressing pYfiN and its I-site double mutant (I_ps_) variant were incubated with [α-^32^P] GTP at regular time intervals from 0-15 min and the products analyzed by thin layer chromatography (TLC); only data for the 10 min time point are shown because in this experiment, WT pYfiN lysates depleted almost all of the input GTP at this time, while ∼20% of input GTP still remained in the I_ps_ mutant (**Fig. 4D**). We then compared the time course of GTP consumption by WT pYfiN with that of I-site regulated pDgcA (**Fig. 4E**). By16 min, YfiN had consumed most of the GTP, while half of the input GTP still remained during DgcA expression. This difference in enzymatic activity may be critical when intracellular GTP is low to begin with in glycerol-fed cells (∼2.5 fold lower compared to glucose) [39]. We finally used a fourth method to verify GTP depletion by quantifying the absolute levels of GTP using mass spec as described under Methods. GTP levels decreased significantly in cells expressing pYfiN (**Fig. 4F**). Although _p_YfiN(I_ps_) and _p_DgcA cells showed higher levels of GTP compared to _p_EmptyVector, the difference was not statistically significant. Thus, four different kinds of measurements show that YfiN expression depletes cellular GTP as expected. The data in Fig. 4B show that YfiN expression impacts all ribonucleotide triphosphate levels as well.

**Figure 4.**
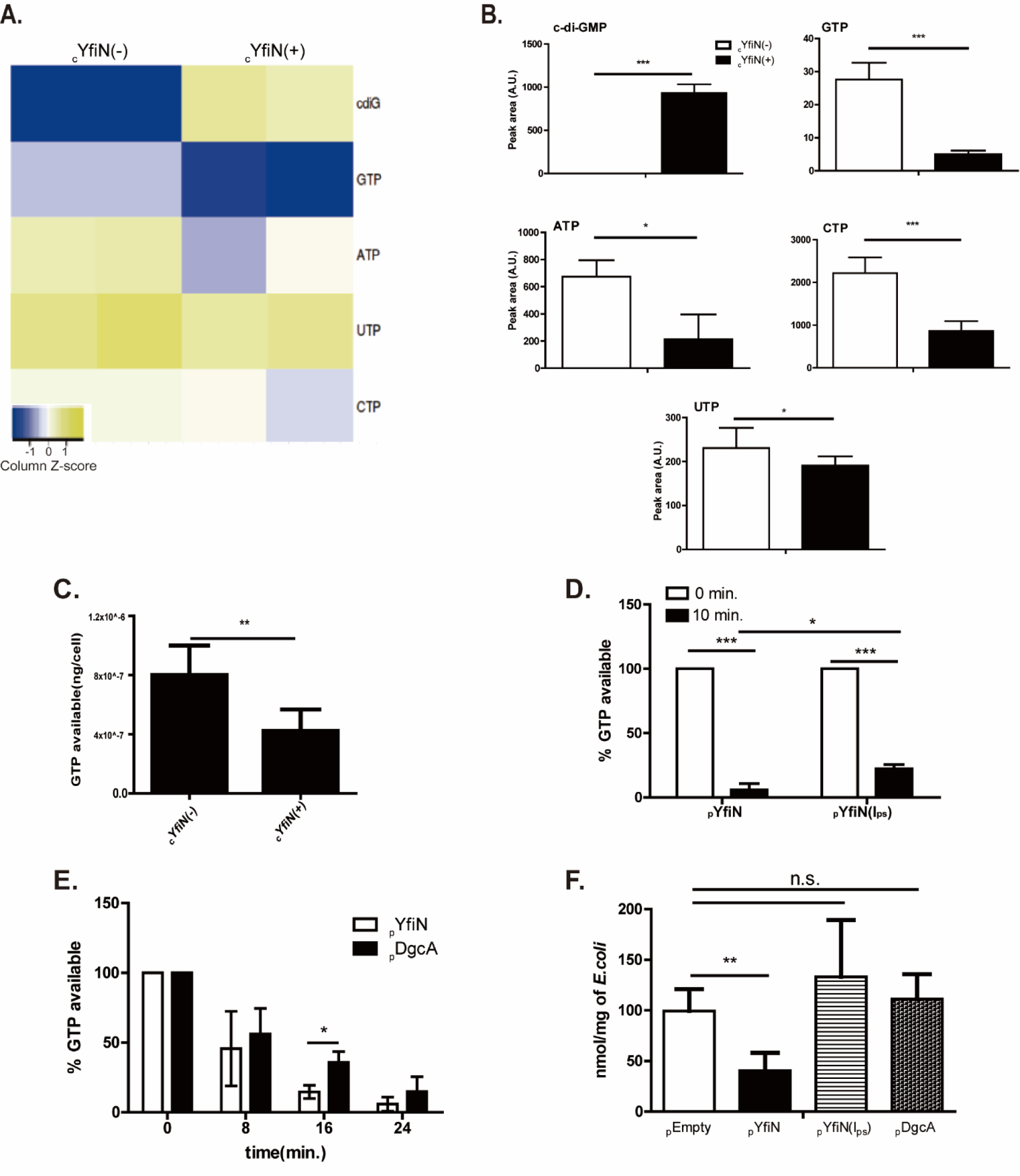
YfiN depletes intracellular GTP. (*A*) Metabolite profiling of cYfINGFP was performed using two biological replicates (each with 2 technical replicates) with and without inducer (+/-) (see Methods). Among ∼219 metabolites measured using the targeted metabolomics approach, levels of c-di-GMP (cdiG) and the four ribonucleotides (GTP, ATP, UTP, CTP) are shown. Peak areas under selected metabolite elution profiles are log10 transformed and displayed as a heatmap, data from each biological replicate (each with two technical replicates) occupying a square. *(B)* To measure the statistical significance of the estimates in A, the areas under the curve of eluted peaks were calibrated as AU (arbitrary units), and the data replotted as bar graphs. (*C*) A luciferase-based GTP assay kit was used to measure GTP levels (ng/cell) in lysates of cYfiNGFP-expressing cells 3 hours post-induction. (*D)* Lysates of M9M-grown cells expressing pYfiN and pYfiN Ips were incubated with [α-32P] GTP for 5-minute intervals from 0-15 minutes, and analyzed using TLC (see Methods); only the 10 min data are shown (see text). Each replicate was normalized to initial GTP levels set at 100%. *(E)* As in *D,* but with lysates of cells expressing pYfiN and pDgcA. *(F)* Absolute quantification of GTP levels in cells expressing indicated plasmids (see Methods). Concentration of GTP for each biological replicate was normalized to wet weight of *E. coli* cells.

An alternative explanation for the depletion of cellular GTP observed in Figure 4 could be activation of the (p)ppGpp synthesis pathway, during which a diphosphate from ATP is transferred to the 3′-OH oxygen of GTP/GDP; (p)ppGpp is the primary regulator of GTP homeostasis in *E. coli* [39], and deployed in bacteria experiencing various stresses [40, 41]. In *E. coli*, RelA and SpoT are the only known enzymes that synthesize as well as degrade (p)ppGpp, which is not made in the absence of both proteins (ppGpp^0^) [42]. To test this alternate explanation, we induced YfiN in *ΔrelA* or *ΔrelAΔspoT*(ppGpp^0^) background, expected to have minimal amounts of (p)ppGpp. Both mutant strains showed growth arrest as early as 2 hours after induction of YfiN (**Fig. S5**), ruling out involvement of (p)ppGpp in YfiN-induced arrest.

To run an independent check of studies concluding that less GTP is available during growth on poor carbon sources such as glycerol and acetate [43], we considered tweaking enzymes contributing to gluconeogenesis. We chose phosphoenolpyruvate carboxy kinase (PCK) for our test, as it mediates an irreversible step in the pathway, converting oxaloacetate to phosphoenolpyruvate, and promotes gluconeogenesis [44] (**Fig. S6A**). Increasing PCK levels might therefore be expected to increase severity of the growth arrest. We monitored growth in M9 mannitol, where cYfiN-mediated growth arrest is not as severe as in M9M (**Fig. 1E**). In this growth medium, PCK induction resulted in a severity of growth arrest similar to that seen in M9M (**Fig. S6B)**. These data, along with those demonstrating that YfiN-mediated growth arrest can be reversed by the addition of a non-gluconeogenic sugar (**Fig. S2E**), support our hypothesis that the availability of GTP pools underlies the mechanism of the YfiN-mediated growth arrest.

In summary, we infer from the data in this and prior sections that absence of feed-back control of c-di-GMP synthesis in YfiN, combined with the energy-expensive metabolism of gluconeogenic sugars, deplete cellular GTP to levels unsustainable for growth, which can be rescued by utilizing non-gluconeogenic sugars.

### YfiN-arrested cells are tolerant to a broad class of antibiotics

To investigate whether all major cellular processes - cell wall, protein and DNA synthesis – are arrested by YfiN during growth on M9M, we exposed cells to the antibiotics Ampicillin, Gentamicin, and Ciprofloxacin [45]. At the bactericidal concentrations used, _c_YfiN-expressing cells survived better than control cells in the presence of all three antibiotics when tested as follows. Cells were first grown with or without cYfiN induction on M9M + antibiotic plates for 12 hours (**Fig. 5A**, middle). They were then transferred with an inoculation loop to LB plates without added antibiotics for recovery (**Fig. 5A**, right; Gen/Amp/Cip stamps only point to the antibiotic condition from where cells were taken). The results are shown in **Figure 5B** (note again that there are no antibiotics in these plates, as illustrated in Fig. 5A, right). Without cYfiN expression, cells succumbed to all three antibiotics they were initially plated on, as judged by loss of recovery of viable cells (**Fig. 5B, left**); this was not the case when cYfiN was expressed (**Fig. 5B, right**). To quantify the number of YfiN-expressing cells that survive antibiotic treatment, a similar assay was performed in M9M, where after 4 hours of antibiotic exposure, survival was measured by CFU counts on LB plates (**Fig. 5C, D**). The results were similar to those reported in Fig. 5B i.e YfiN-expression increases survival to multiple antibiotics. We conclude that YfiN effectively shuts down all major cellular processes in M9M.

**Figure 5.**
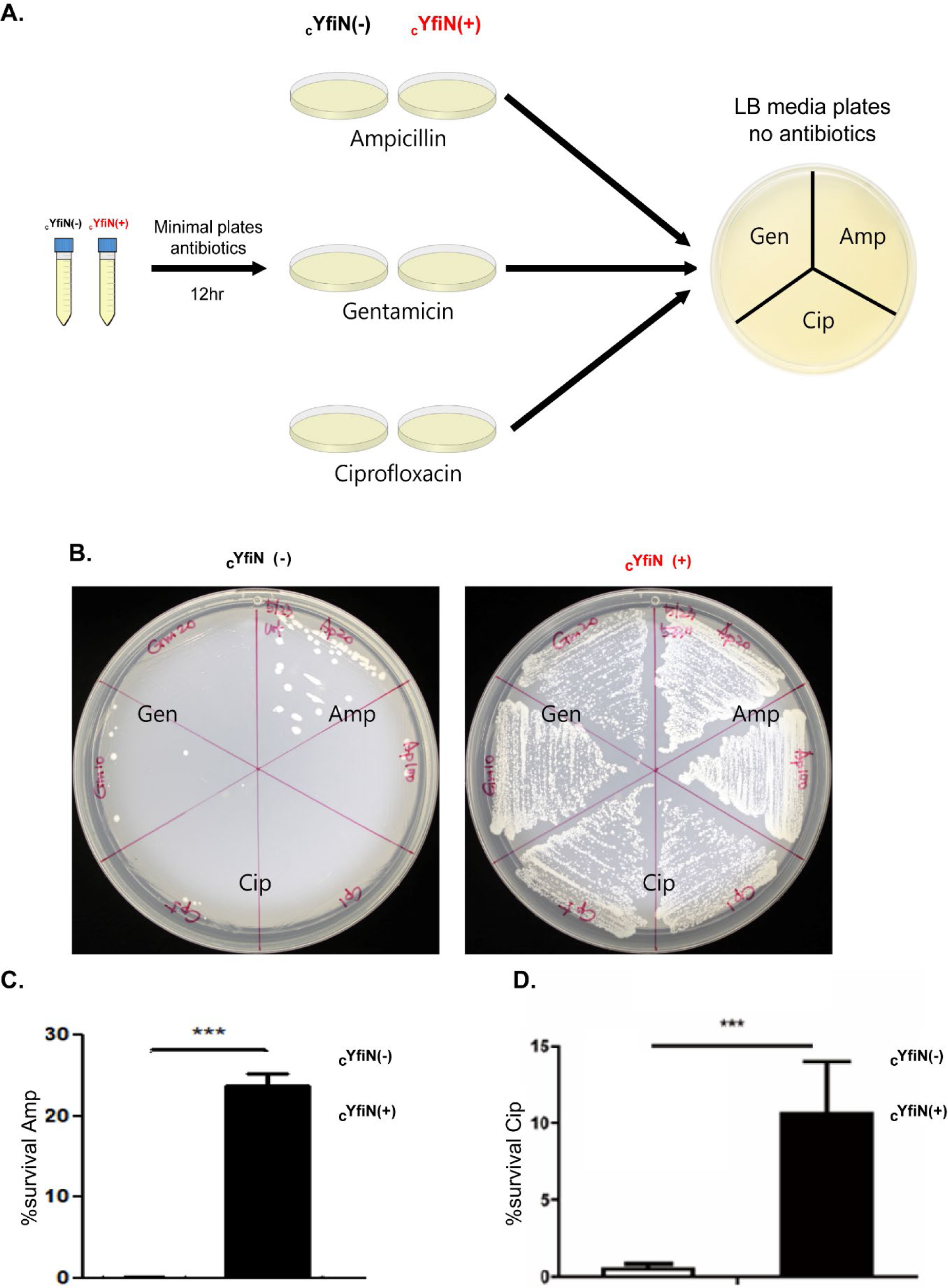
YfiN increases tolerance to different classes of antibiotics. *(A)* Experimental design. (cYfiN**GFP**)-expressing cells grown with or without inducer (+/-) in liquid M9M media were plated on M9M plates supplemented with two different antibiotic concentrations: Ampicillin (Amp; 20 & 100ug/ml), Gentamicin (Gen; 4 & 20ug/ml), Ciprofloxacin (Cip; 2 & 10ug/ml). After overnight incubation, inoculating loops were used to transfer cells to fresh LB plates without antibiotics. The results are shown in *(B)*. Note that there are no antibiotics in the plates shown in B; the labels merely indicate the antibiotic they were treated with on the M9 plates prior to inoculation on the LB plates. *(C-D)* In a separate set of experiments, cells grown with or without YfiN (cYfiN**GFP**) in M9M liquid media for 3 hours were exposed to (*C*) Amp (100ug/ml) or (*D*) Cip (10ug/ml) for 4 hours, before plating on LB. % survival under the three conditions was determined by measuring CFU counts before and after exposure to antibiotics.

Since the enzymatic function of YfiN is closely related to the growth arrest phenotype, we reasoned that this function should also be linked to its antibiotic tolerance property. This was tested in two ways using _c_YfiN – by concomitantly increasing the expression of its periplasmic inhibitor YfiR (pYfiR) (**Fig. S7A**), as well as by testing the GGDEF active site mutant (**Fig. S7B**). Both manipulations abrogated the antibiotic (Cip) tolerance phenotype. We conclude that the DGC activity of YfiN mediates both growth arrest and antibiotic tolerance.

### Native YfiN delays exit from lag phase during exponential growth

The results presented thus far have relied on ectopic expression of YfiN from non-native promoters, either from the chromosome or from a plasmid. But are they relevant to the native situation? Since we do not know what environmental conditions induce native YfiN, we took advantage of a report that showed a brief burst of YfiN protein levels when overnight cultures are inoculated into fresh media [46]. We therefore prepared overnight cultures of WT and *ΔyfiN* strains in LB, and inoculated them to either M9 glycerol or M9 glucose, recording their growth rates over three 2-hour intervals are shown in **Fig. 6AB** (*ΔyfiN* data are in orange). In M9 glycerol (**Fig. 6A**), both strains were still in the lag phase in the first growth interval (0-2 hour). In the next interval (2-4 hour), the growth rate of WT was unchanged i.e. it was still in the lag phase, but that of the *ΔyfiN* strain increased. In the third interval or exponential phase of growth (4-6 hour), both strains grew at the same rate. Thus, the WT strain showed a longer lag or delayed transition to exponential phase compared to *ΔyfiN.* In M9 glucose, however, both strains grew similarly, and the early lag seen in M9 glycerol for WT was not observed (**Fig. 6B**). To test whether this observation was strain specific, we repeated the same experiment using WT and isogenic *ΔyfiN* derivatives of the uropathogenic *E. coli* strain CFT073. The longer lag pattern seen for our WT strain (MG1655) in glycerol (**Fig. 6A**), was also seen for CFT073 compared to its *ΔyfiN* counterpart (**Fig. S8**).

**Figure 6.**
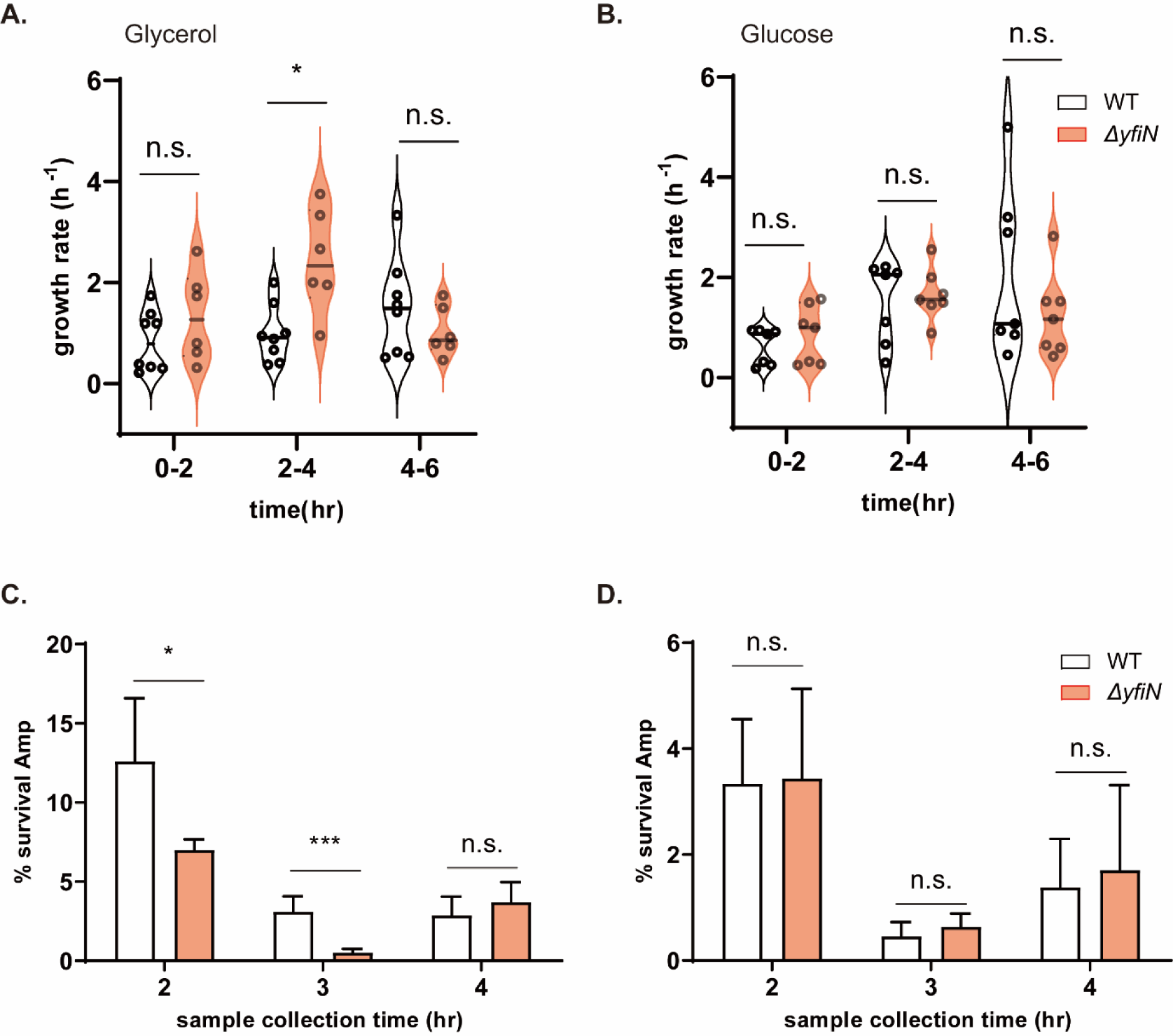
Native YfiN increases the duration of the lag phase of growth. (*A-B*) WT versus *ΔyfiN* strains were grown overnight in LB and inoculated in either M9 glycerol or glucose. Growth rates in the M9 media were calculated using CFUs measured at each time interval shown (see Growth Rate Calculation under Methods). Each black circle is a biological replicate (n=8 glycerol, n=7 glucose). (*C*-*D*) WT and *ΔyfiN* strains were grown in M9 glycerol or glucose for 0-4 hours and aliquots removed at 2, 3, and 4 hours (within the time windows monitored in A), and treated with Ampicillin (100μg/ml) for 4 hours. At each time, % survival was calculated by comparing CFU counts before and after antibiotic treatment.

To test if the antibiotic tolerance seen during growth delay/arrest by ectopic expression of YfiN (**Fig. 5**) would also be seen under native conditions where growth of the WT strain was delayed (**Fig. 6A**), we sampled aliquots of the YfN +/- strains at 2, 3, and 4 hour (which fall within the time windows monitored in **Fig. 6A/B**), and measured their ability to survive a 4-hour exposure to Ampicillin. The survival data for the WT strain inoculated in either glycerol for glucose could be superimposed on growth periods that coincided with the extended lag for the WT strain (**Fig. 6AB**) i.e. WT survived better than Δ*yfiN* after antibiotic treatment only during the lag phase and only in M9M (compare **Fig. 6C** and **Fig. 6D).** The larger killing effect of Ampicillin on the strains grown on glucose could be due to the reported enhanced killing on this metabolite [47, 48].

If the lag was due to a burst of YfiN synthesis as reported [46], this should be reflected in c-di-GMP levels. These were monitored by the riboswitch sensor at the same time points where antibiotic survival was measured i.e. 2, 3 and 4 hour. The data are plotted as a ratio of c-di-GMP in WT vs Δ*yfiN* (**Fig. S9**). Compared to Δ*yfiN,* the WT strain shows higher c-di-GMP levels at 2 and 3 hour but not at 4 hour, mirroring the antibiotic survival pattern in **Fig. 6C**.

In summary, the growth delay/arrest property of YfiN is seen even when the protein is expressed from its native chromosomal location, and is not an artifact of ectopic expression.

## Discussion

C-di-GMP is the most ubiquitous signaling nucleotide in bacteria, with dozens of DGCs involved in its production. These enzymes enable a variety of downstream outputs. In this study, we have discovered a new output for the DGC YfiN, made possible by a particular structural feature of the enzyme that restricts growth in specific nutrient conditions, allowing *E. coli* to survive through stressors like antibiotics.

### YfiN exploits the absence of autoinhibitory I sites to enable a novel mode of survival

I sites (I_p_-I_s_) are common in DGCs, and serve an important role in binding c-di-GMP to feed-back regulate DGC activity [12, 20], thus controlling the amount of c-di-GMP available to bind to downstream effector proteins. YfiN belongs to a small fraction of DGCs in *E. coli* (3/12, based on sequence gazing) that do not encode these sites. Absence of this regulatory structural feature appears to be the hallmark of the majority (>90%) of all bacterial YfiN homologs found in the UniProt database. YfiN activity is instead controlled by the periplasmic repressor YfiR, known to be inactivated by redox stress [16, 24]. Given that YfiN is the most robust DGC in *E. coli* [24], we suspect other periplasmic stresses that unfold proteins may also activate this enzyme [49]. Signals that might activate transcription of *yfiN* are still unknown. Once activated, one would not expect YfiN to stop c-di-GMP synthesis until the inducing stressors are gone. We show in this study that unregulated c-di-GMP production comes with a metabolic cost. The nature of this cost came to the fore when *E. coli* grown on gluconeogenic carbon sources such as glycerol, mannitol, or sorbitol were observed to arrest cell growth, a phenotype that required the DGC activity of YfiN (**Fig. 1**), but did not require its relocation to the cell division site at the mid-cell (**Fig. 2**). That it was the unregulated DGC activity of YfiN that was responsible for growth arrest was established by reconstituting the consensus sequence of both I_p_ and I_s_ sites, which restored growth (**Fig. 3C**). Conversely, inactivating the I_p_ site of DgcA from *C. cresentus*, conferred on it the growth arrest phenotype (**Fig. 3E**).

The studies described above were performed under ectopic expression of YfiN alone, in order to mimic conditions where YfiR is non-functional. Deleting *yfiR* was not sufficient to induce growth arrest from the native levels of YfiN. While we do not as yet know which environmental signals might activate *yfiN* expression, a brief spike in YfiN levels was reported when an overnight culture of *E. coli* was inoculated into fresh media [46]. As discussed below, we have leveraged this finding to show that during this brief spike, native YfiN recapitulates the data from ectopic expression. We imagine that in natural habitats, a combination of periplasmic stress plus the stress of growing on energetically costly substrates, combined perhaps with environmental signals that activate *yfiN* transcription, might create fertile grounds for growth arrest. That this arrest is reversible, as seen by revival of growth upon adding non-gluconeogenic sugars (**Fig. S2E**), suggests that *E. coli* employs YfiN to weather inhospitable conditions. A reversible quiescent state is known to favor adaptive evolution from microbes to humans [50–52]. Our findings likely extend to all YfiN-encoding bacteria given that the majority of the homologs lack the I_p_ site. In those that do have this site (*Pseudomonas* and *Yersinia* species), the I_s_ site is absent (**Fig. S10**). In light of our data showing that both I sites are required to relieve growth arrest (**Fig. 3C**), feed-back inhibition of YfiN activity is likely inefficient in the bacteria harboring only the I_p_ site [21, 22].

### YfiN activity during growth on gluconeogenic carbon sources depletes cellular GTP

The clear distinction between gluconeogenic and glycolytic carbon sources in promoting growth arrest by YfiN is striking (**Fig. 1D**). Gluconeogenesis is the process by which cells synthesize glucose 6-phosphate from non-hexose sugars [53]. While eukaryotic cells as well as other bacteria consume both ATP and GTP in this process, *E. coli* is known to only consume ATP [54]. Each cycle of gluconeogenesis requires 4 ATPs plus 2 NADHs, resulting in net deficit of 9 ATPs. Lower levels of ATP will consequently lower those of GTP, since the γ phosphate of ATP is used by nucleoside diphosphate kinase (ndk) to synthesize GTP from GDP [55] (**Fig. 7**). In glycerol, there is less ATP produced during TCA cycle as well (20 ATPs compared to 36∼38 ATPs in glucose). Indeed, *E. coli* grown in M9 glycerol and acetate has been demonstrated to have less intracellular ATP/GTP than in glucose [43]. Unregulated consumption of GTP by YfiN will further increase cellular metabolic stress under these conditions. The lower cellular GTP levels we see upon induction of YfiN during growth on M9M are therefore expected (**Fig. 4A**). Lowering GTP levels even by two-fold (**Fig. 4B,C,F**) should be sufficient to arrest growth given *in vitro* data showing that the transcription rate from a ribosomal RNA promoter (*rrnB*) is highly sensitive to GTP concentrations [56]. It was reported earlier that depletion of intracellular GTP by 50% along with decrease in ATP levels, by overexpression of RelA homologues contributes to persister formation by arresting cell growth [57]. In eukaryotes, cancer drugs that cause similar drop in GTP levels resulted in cell death in a human cancer cell line [58]. In the ecological niches that heterotrophic bacteria inhabit, they obtain carbon from dissolved organic matter. Although *E. coli* is primarily a commensal of mammals, and to a lesser extent birds, it can be isolated from a variety of host species as well as soil, sediments, and water. We imagine that the particular YfiN function we have uncovered in this study may manifest in certain ecological niches rich in gluconeogenic sugars or in non-carbohydrate substrates not tested in this study. For example, each part of the human gut has different concentrations of carbon sources and types [59].

**Figure 7.**
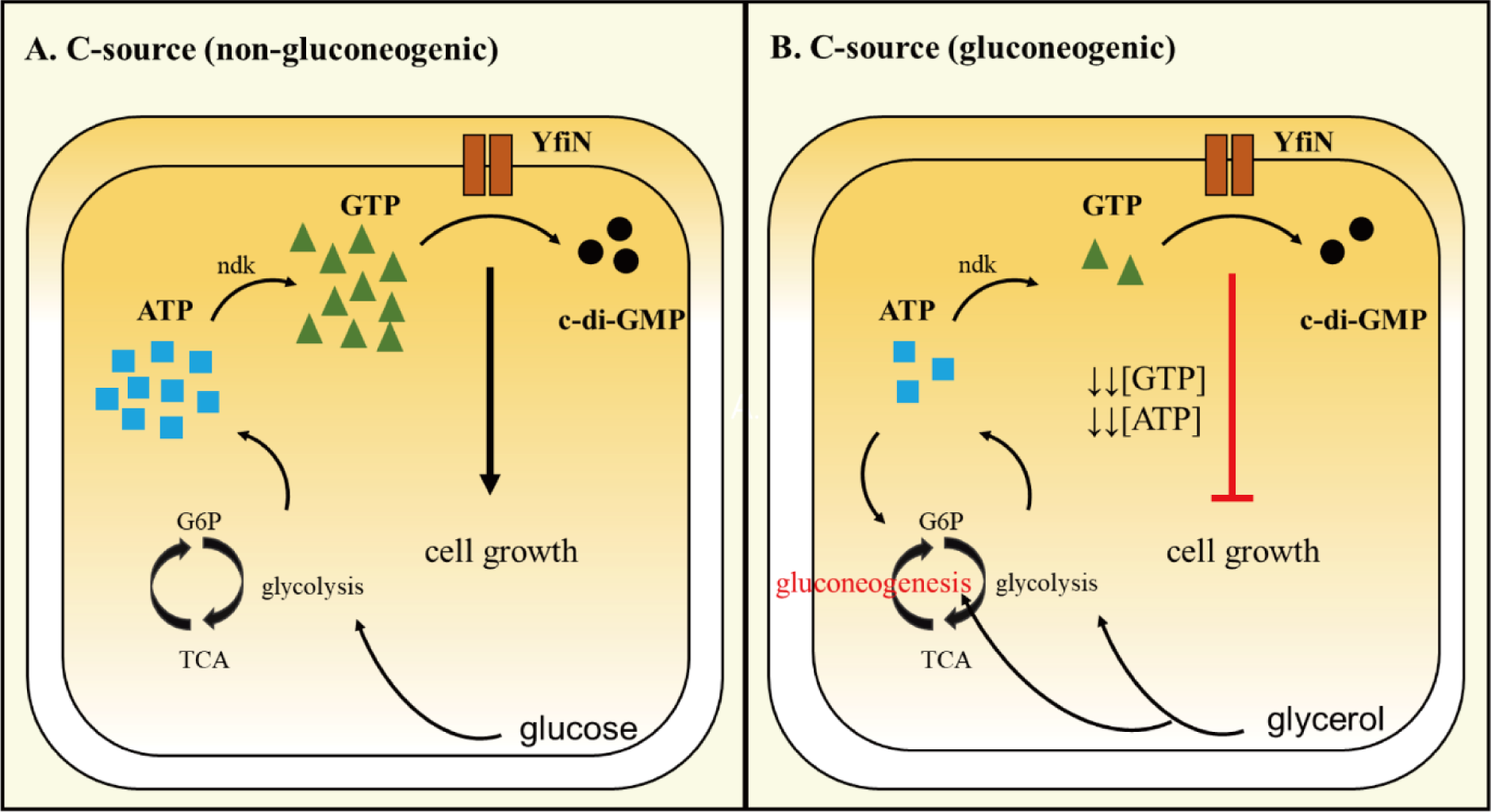
Model for mechanism of YfiN-mediated growth arrest on gluconeogenic carbon (C) sources. (*A*) On glycolytic substrates or rich media, GTP consumption by YfiN does not deplete NTP pools because they are abundant. (*B*) On non-gluconeogenic carbon sources NTP pools are low to begin with, and easily depleted by YfiN activity.

In summary, we conclude that uncontrolled synthesis of c-di-GMP by YfiN is the proximate cause of depletion of cellular GTP. These nucleotides are essential for the synthesis of the major cellular macromolecules DNA, RNA and protein, explaining why cells enter growth arrest when using the energetically more costly gluconeogenic carbon sources for synthesis of glucose-6-P.

### YfiN contributes to increased tolerance to antibiotics

YfiN is known to inhibit cell division in response to redox and envelope stresses, and to protect *E. coli* from envelope-disrupting environments when ectopically expressed [24]. We show here that ectopic expression of YfiN in M9M also increases tolerance to a broad class of antibiotics such as Ampicillin, Gentamicin, and Ciprofloxacin, which target and disrupt a variety of cellular processes (**Fig. 5**). This is a result of YfiN’s unregulated DGC activity (**Fig. 1C**, **3C**). Bacteria are continuously exposed to environmental stresses and knowing when to stop proliferating is key to their survival. Our finding that transferring *E. coli* cells from a stationary culture to M9 glycerol delays growth by extending the lag phase (**Fig. 6AB**), might suggest that transitioning from one nutritional environment to another is one such stress that activates YfiN expression [46]. The growth delay upon transitioning from LB to M9M is accompanied by increased c-di-GMP levels (**Fig. S9**), and increased tolerance to antibiotics compared to a Δ*yfiN* strain (**Fig. 6C**). This result is similar to antibiotic tolerance observed when cell growth is delayed/arrested by ectopic expression of YfIN. Although it may seem counter-intuitive to harbor a gene preventing bacteria from proliferating, persisting in the lag phase may help bacteria in transitioning to the new environment, particularly if the environment is stressful or nutrients are scarce. An increase in the lag phase has been reported as an integral step for developing antibiotic resistance [60, 61]. It has been suggested that a prolonged lag could buy the bacteria time to diversify adaptive phenotypes [62, 63]. The importance of the lag phase is not limited to surviving antibiotic stress alone. A transcriptional profiling study of *S. enterica* showed that genes associated with DNA repair and protein degradation were induced during lag phase, implicating a crucial role for this phase in repairing damaged cellular components [61, 64]. Bacteria are also known to reorganize their metabolism during the lag phase to achieve optimal growth [64], and two distinct lag phases were observed in *E. coli* supplied with arabinose [65]. Thus, by regulating the duration of the lag phase, we suspect that YfiN not only confers increased tolerance to antibiotics but also other advantages such as optimizing DNA repair and metabolic pathways for growth.

#### Coda

YfiN seems to have emerged as a DGC that responds to multiple metabolic stresses – redox, envelope, gluconeogenic substrates – and has thus far shown multiple output responses designed to hunker down, make biofilms, and persist.

## Acknowledgements

We thank Y. Fitnat, HK. Kim and S. Bhattacharyya for strains and for helpful discussions. This work was supported by NIH grant R35GM118085.

## Materials and Methods

### Strains, growth conditions, mutagenesis, and plasmid constructions

Strains and plasmids used in this study are listed in Table S1. The WT parent strain for *E. coli* was MG1655 for all experiments. All strains were grown in M9M (M9 minimal media + 0.2% glycerol + 0.2% casamino acid) or in LB broth (10 g/liter tryptone, 5 g/liter yeast extract, 5 g/liter NaCl) unless noted otherwise. When appropriate, the following antibiotics were used: Ampicillin (100 µg/ml), Chloramphenicol (20 µg/ml), Kanamycin (50 µg/ml), and Gentamicin (30 µg/ml). For inducible plasmids, 100μM isopropyl-β-d-thiogalactopyranoside (IPTG) or 0.02%(w/v) L-arabinose were added as indicated in the figures or legends. To monitor the growth of cells, optical density was measured at the wavelength of 600nm (OD_600_)

HK 533 strain was constructed similarly to HK532 strain [24]. Using lambda red recombination [66], *yfiR* was replaced with PTrc promoter and a kanamycin cassette inserted in an orientation opposite to the direction (←) of *yfi* operon transcription, in order to prevent polar effects on *yfiN* expression.

For cloning of expression plasmids, gene sequences were amplified from the genomic DNA of WT strains by using PCR and introduced into pBAD30, pBAD33.

To restore or introduce I-sites, specific primers designed for single base pair substitutions were used for PCR amplification using pBAD_YfiN or DgcA as templates. Following amplification, the original templates were digested with 1 unit of DpnI and PCR products were used for transformation and selection. All constructs were confirmed by DNA sequencing.

### Recovery assay

For the experiment shown in Fig. S2E, cYfiN strain was propagated in M9M with IPTG for induction. After 2 hours post-induction, 0.2% glucose was added for recovery, followed by measurement of OD_600_ every 2 hours.

### Motility assay

LB soft agar or swim plates were made using 0.3% Fisher agar. 5 µl of an overnight culture was inoculated in the center and plates incubated at 30°C for 8-12 hours. Swim ring diameter was measured to compare motility across strains.

### Biofilm assay

Cells were propagated in M9M in 96-well plates. After 20-hour incubation at 37℃, plates were washed twice with water and dried for 2 hours at RT. 125μl of 0.01% Crystal violet solution (w/v) was added to each well and incubated in RT for 15 min. The plate was re-washed three times with water and then 125μl 30% Acetic acid (v/v) were added to each well.

After 15 min incubation in RT, solutions in each well were transferred to a new 96 well plate. O.D. 550nm was used to measure biofilm formation, and OD_600_ for data normalization.

### Fluorescence microscopy

Overnight cultures of cells with plasmids encoding fluorescent fusion proteins were diluted 1:100 in fresh M9M or LB medium with antibiotics and grown at 30°C with 0.02% arabinose for 4 hours (unless otherwise stated). For imaging cells, a cell suspension (5 µl) was applied to a slide and incubated for 5 min before imaging. Images were acquired using an Olympus BX53 microscope, appropriate filters, and cellSens standard software (version 1.6) from Olympus.

### Western Blot

Overnight cultures of cells with plasmids encoding YfiN and its mutants (I_ps_, R260A, GGAAF) were diluted 1:100 in M9M and propagated with 0.02% arabinose for 3 hours. Cells were collected and then re-suspended in a lysis buffer with a final concentration of 2.5×10^9/ml cells. Upon cell lysis using a heat block (100℃), 1×10^7 cells were loaded into each lane on a SDS-gel. Proteins were then transferred to a nitrocellulose membrane and blocked with 5% <u>n</u>on-fat <u>d</u>ry <u>m</u>ilk (NFDM) in <u>T</u>ris-<u>b</u>uffered <u>s</u>aline with 20% <u>T</u>ween (TBST) for 1 hour. The membrane was incubated overnight with a 1:1000 dilution of either GFP antibody (Sigma) or FLAG antibody (Sigma Monoclonal FLAG antibody M2) in TBST with 5% NFDM (w/v), and then washed 3 times with TBST every 10 minute intervals. This step was followed immediately by an hour incubation with 1:5000 dilution of Goat α-mouse-HRP (Bio-Rad) in TBST. ECL Select (Amersham; chemiluminescence) was used for a development and the blot image was taken using ChemiDoc (G:BOX).

ImageJ software was used to measure the difference in pixel intensities to compare the difference in the levels of expression.

### c-di-GMP biosensor assay

Overnight cultures of cells with plasmids encoding YfiN and riboswitch-based c-di-GMP biosensor ere diluted (1:100) and then propagated in M9M media with an addition of 0.02% arabinose for induction. After a 3-hour incubation, cells were transferred to a 96-well plate (100μl/well). Riboswitches (bc3,4,5) that are situated upstream of *turborfp* (encoding a more photo-stable RFP variant) are activated upon binding of c-di-GMP and allow for downstream gene expression, while *amcyan,* an enhanced CFP variant, is expressed constitutively. Samples were diluted to OD_600_ prior to measurement and the ratio of RFP/CFP was used to estimate c-di-GMP concentration/cell. Excitation/Emission wavelengths of 405/488nm and 553/574nm were used for *amcyan* and *turborfp* respectively using a FlexStation3 Plate Reader.

### Gene alignment for identification of primary and secondary I-sites

Primary I-site designation for YfiN was based on the identified consensus RxxD, situated five amino acids upstream of the GGDEF catalytic site [12, 19, 67]. Secondary I-site designation for YfiN was predicted by comparing sequences from other DGCs that harbor experimentally identified secondary sites. Clustal Omega, available online (https://www.ebi.ac.uk/), was used for the prediction and alignment; DgcA was excluded from the analysis, because it has two predicted secondary I-sites.

For analysis of YfiN homologues, we used YfiN protein sequence of *E. coli* MG1655 from the UnitProt database (http://www.uniprot.org; accession number: P46139). Using UniRef50, we acquired 1974 orthologous sequences with minimum 50% sequence identity. Conservation of primary and secondary I-sites was calculated using the Clustal Omega tool.

### Antibiotic survival assay on plates

An overnight culture of cYfiN strain was diluted 1:100 in M9M with or without IPTG. After 6 hours post-induction, cells were spread on agar plates containing either Ampicillin (20, 100 µg/ml), Gentamicin (10, 20 µg/ml) or Ciprofloxacin (1, 5µg/ml). After 12 hour incubation on these plates, cells were re-spread on fresh LB agar plates without antibiotics.

### Antibiotic survival assay in liquid

An overnight culture of cYfiN strain was inoculated at 1:100 dilution in M9M. Cell cultures were grown for 2 hours to reach the OD_600_ of 0.1, at which IPTG was added. Two hours post-induction, either Gentamicin (20 µg/ml) or Ciprofloxacin (10µg/ml) was added. After a 4 hour treatment with antibiotics, cell cultures were washed with PBS (phosphate buffered saline) buffer twice and plated on LB agar for CFU counts. The percent survival was calculated as follows: (final CFU/CFU before antibiotic treatment) × 100. The results are presented as the average results from at least 3 biological replicates.

### Thin layer chromatography assay

YfiN, YfiN I-site mutants (I_p_, I_s_, I_ps_), and DgcA were induced from pBAD plasmids with arabinose in M9M for 3 hours. Then, cells were washed twice with PBS buffer and lysed with a VCX-750 Vibra-Cell sonicator at 20kHz (30 sec pulse followed by 10 sec cool-down for total of 5 cycles). Cell lysates were incubated with 1 mM GTP/[α-^32^P]GTP (0.1 μCi/μl) (PerkinElmer) and incubated at 30°C for various time intervals for comparison pYfiN and pYfiN(Ips). Reactions were stopped by the addition of 1 volume of 0.5 M EDTA. Radio-labeled products were analyzed by polyethyleneimine-cellulose thin-layer chromatography (TLC; Millipore), by spotting 2 μl samples onto TLC plates. Plates were dried at RT for 5 min, and developed in 1.3 M KH_2_PO_4_ (pH 3.2) [12]. TLC data was analyzed with GE Typhoon Phosphorimager and ImageJ software by comparing the intensities of each blots on the TLC plate. GTP consumption was estimated based on the input GTP concentration.

### GTPase assay

The GTP levels were measured using a GTPase-Glo Assay kit (Promega) which employs an ATP-linked luciferase reaction. Total concentration of GTP in cells was calculated based on the standard curve generated using a standard provided by manufacturer, and normalized to OD_600_ (OD_600_ 1.0 = 5×10^8 cells). The final concentration was represented in ng/cell.

### Metabolomics

*E. coli* MG1655 yfiR::kan pTrc99a-YfiN_GFP_ (HK533) was grown at 30°C for 2 hours in 10 ml M9M, with IPTG for induction. Cells were incubated for additional 3 hours post-induction (OD_600_ reached around 0.4). Pellets were collected and flash frozen. They were shipped on dry ice to the Metabolomics Core Facility at the Mayo Clinic at Rochester, which routinely analyzes a panel of 219 commonly investigated metabolites such as dNTPs, NTPs, carbohydrates, and acids. We provided them with c-di-GMP, and they estimated these concentrations [68]. Samples were analyzed by high-performance liquid chromatography– tandem mass spectrometry (HPLC-MS/MS) at the facility, using a Thermo Fisher Q Exactive mass spectrometer. The analysis is qualitative, and data are provided as areas under elution peaks (AU, arbitrary unit). Of the 3 biological replicates (each with two technical replicate) sent to the Facility, one was unusable. COVID-19 work protocols during this time (August to November, 2020), prevented us from sending more samples.

Absolute quantification of the targeted nucleotides c-di-GMP and rNTPs was done by (HPLC-MS/MS) at the Metabolomics Core Facility at the University of Texas Medical Branch (UTMB). Cells were grown in M9M except with arabinose for induction. After 3.5 hours post-induction, cells were collected and their wet weight was measured for data normalization. Pellets were then flash frozen and mailed to UTMB. Three biological replicates for each set consisting _p_Empty, _p_YfiN, _p_YfiN(I_ps_), _p_DgcA were prepared. The facility used ^13^C-labeled standards to obtain a standard curve against which experimental values were derived.

### Growth rate calculation

Strains grown in M9M were collected at 0, 2, 4, and 6 hours for counting CFUs. Growth rate h^-1^ = (CFU_final_/CFU_initial_)/2.

## Supplementary Figures and Tables

**Figure S1.**
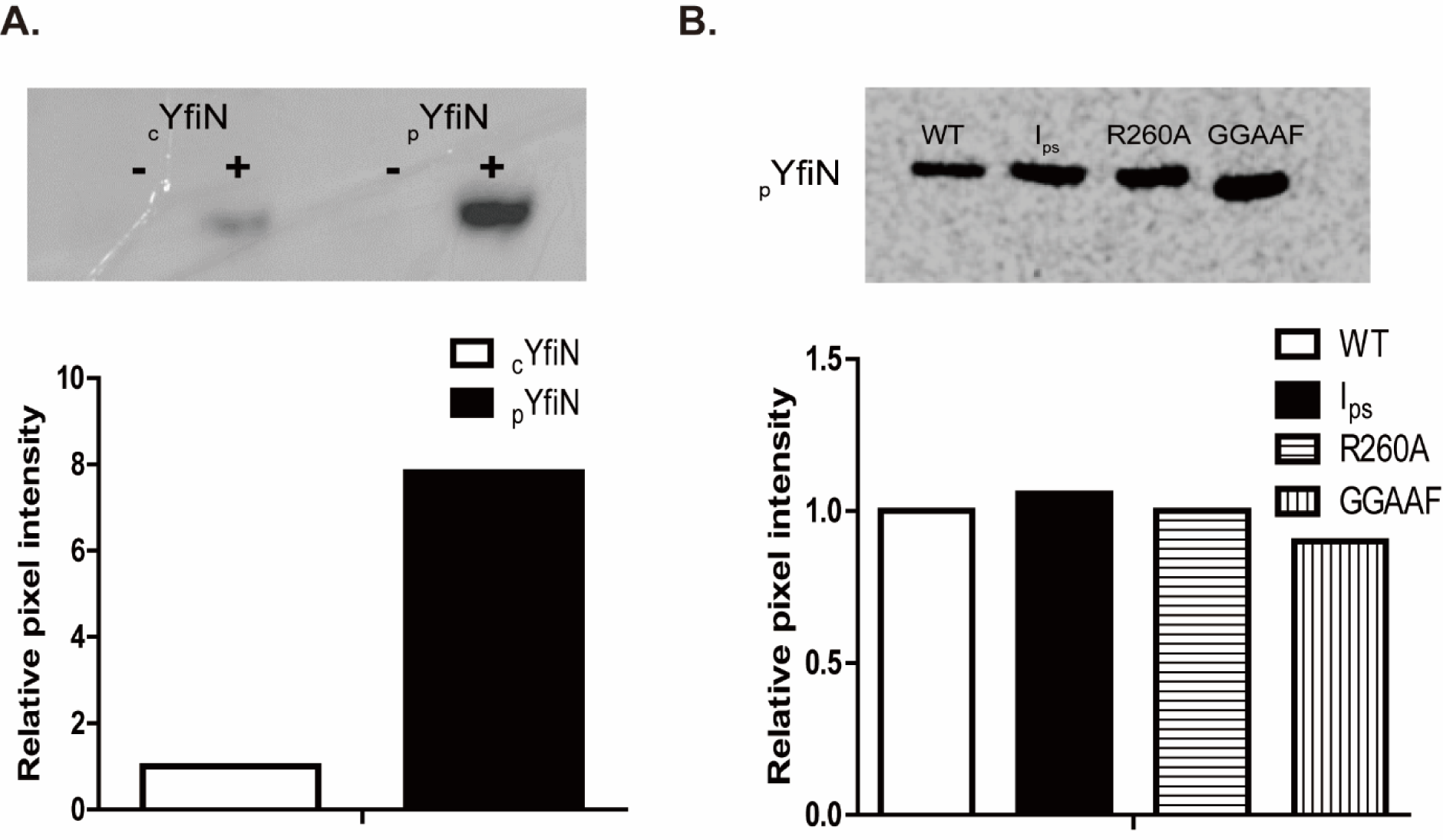
Comparison of protein expression in different YfiN constructs. (*A*) A western blot image of cells expressing cYfiN-or pYfiN-GFP, propagated in M9M with 0.02% arabinose. The image was taken using ChemiDoc and pixel intensities (plotted below) were measured with ImageJ software (see Methods). Intensities were normalized to cYfiN. (*B*) As in *A*, except using Flag-tagged pYfiN(WT), pYfiN(Ips), pYfiN(R260A), and pYfiN(GGAAF), with intensities normalized to pYfiN(WT). The intensities are an average from two replicates.

**Figure S2.**
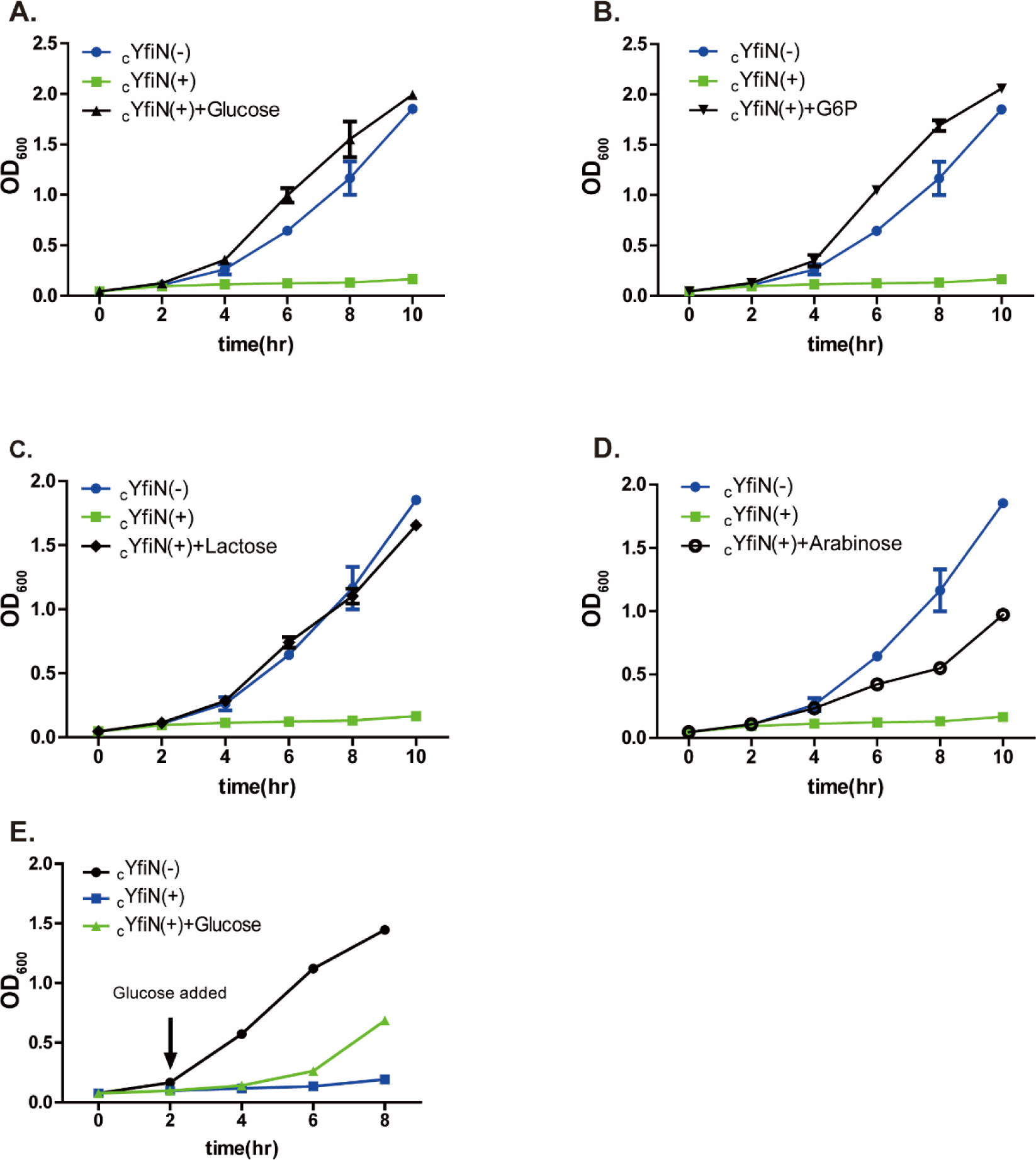
Glycolytic sugars rescue growth-arrest caused by gluconeogenic sugars. *(A-D)* Cells expressing cYfiNGFP were propagated in glycerol alone or in combination with glycolytic substrates (glycerol is present in all experiments, hence not indicated) *(E)* The cYfiN strain was propagated in M9M, and glucose was added at 2 hours post induction. All substrates were at 0.2% w/v. Induction conditions as in Fig. 1.

**Figure S3.**
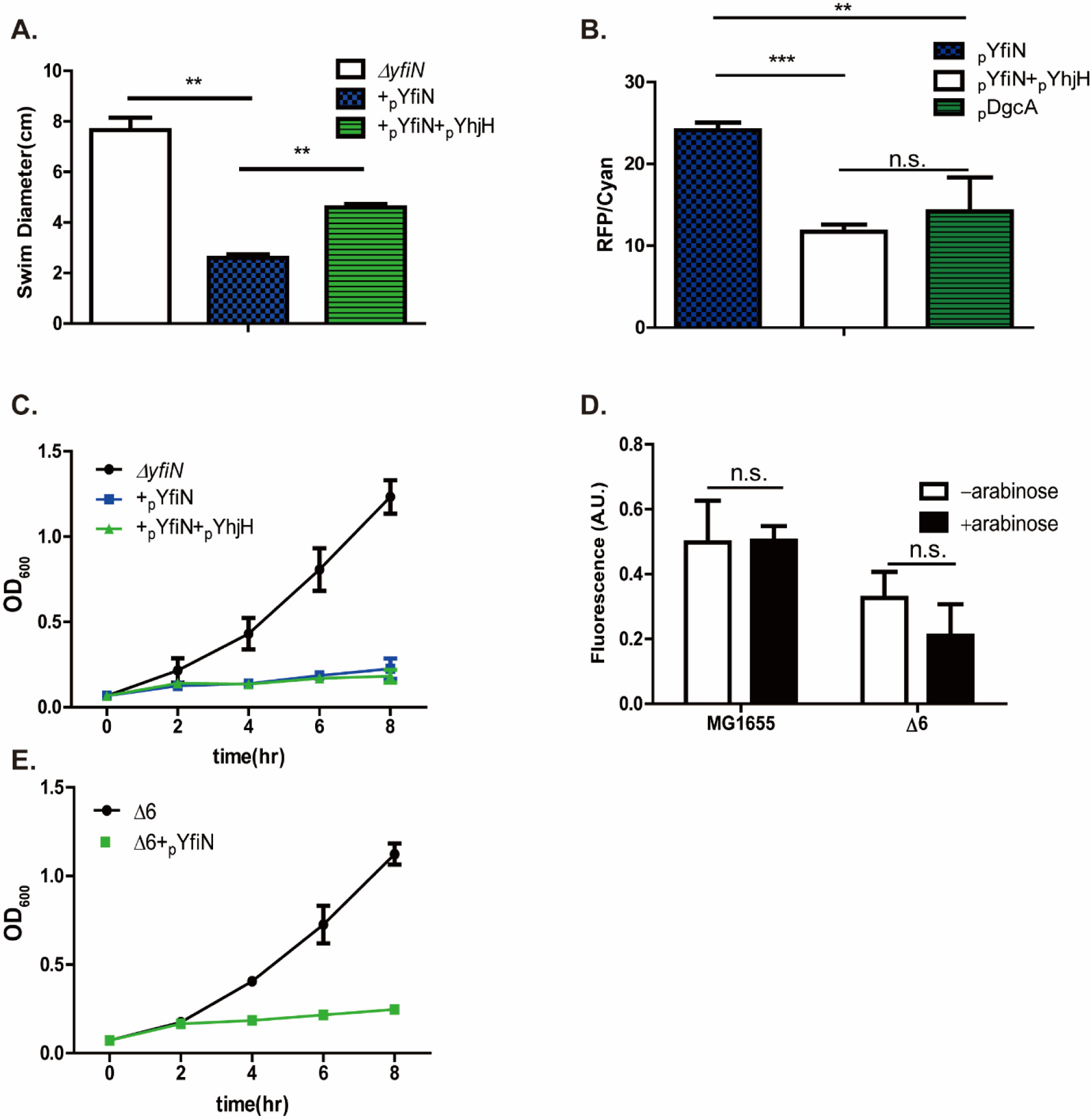
The growth-arrest phenotype of YfiN is unrelated to its motility or biofilm output. (*A*) pYfiN or pYfiN+pYhjH were introduced into the Δ*yfiN* strain and expression induced with arabinose, or arabinose+IPTG, respectively. Motility was measured by estimating the diameter of the swim colony in soft agar plates. *(B)* c-di-GMP levels for pYfiN, pYfiN + pYhjH, and pDgcA were measured using a riboswitch-based fluorescent biosensor, where CFP is expressed constitutively and RFP is the readout for c-di-GMP; the RFP/CFP ratio estimates c-di-GMP/cell (see Methods). The data were obtained from four biological replicates of each sample. (*C*) Strains in *A* were monitored for growth in M9M in the presence of their respective inducers. *D*) The parent MG1655 strain was deleted for six genes that control all the biofilm pathways in *E. coli* (Δ6 = *bcsA, csgD, pgaC, fimA, wcaD, yjbE*). pYfiN was introduced in both strains and the amount of biofilm formed was estimated by Crystal violet staining, with or without arabinose in 96 well plates. A.U., arbitrary fluorescence units. (*E*) Strains in *D* were monitored for growth in M9M in the presence of arabinose.

**Figure S4.**
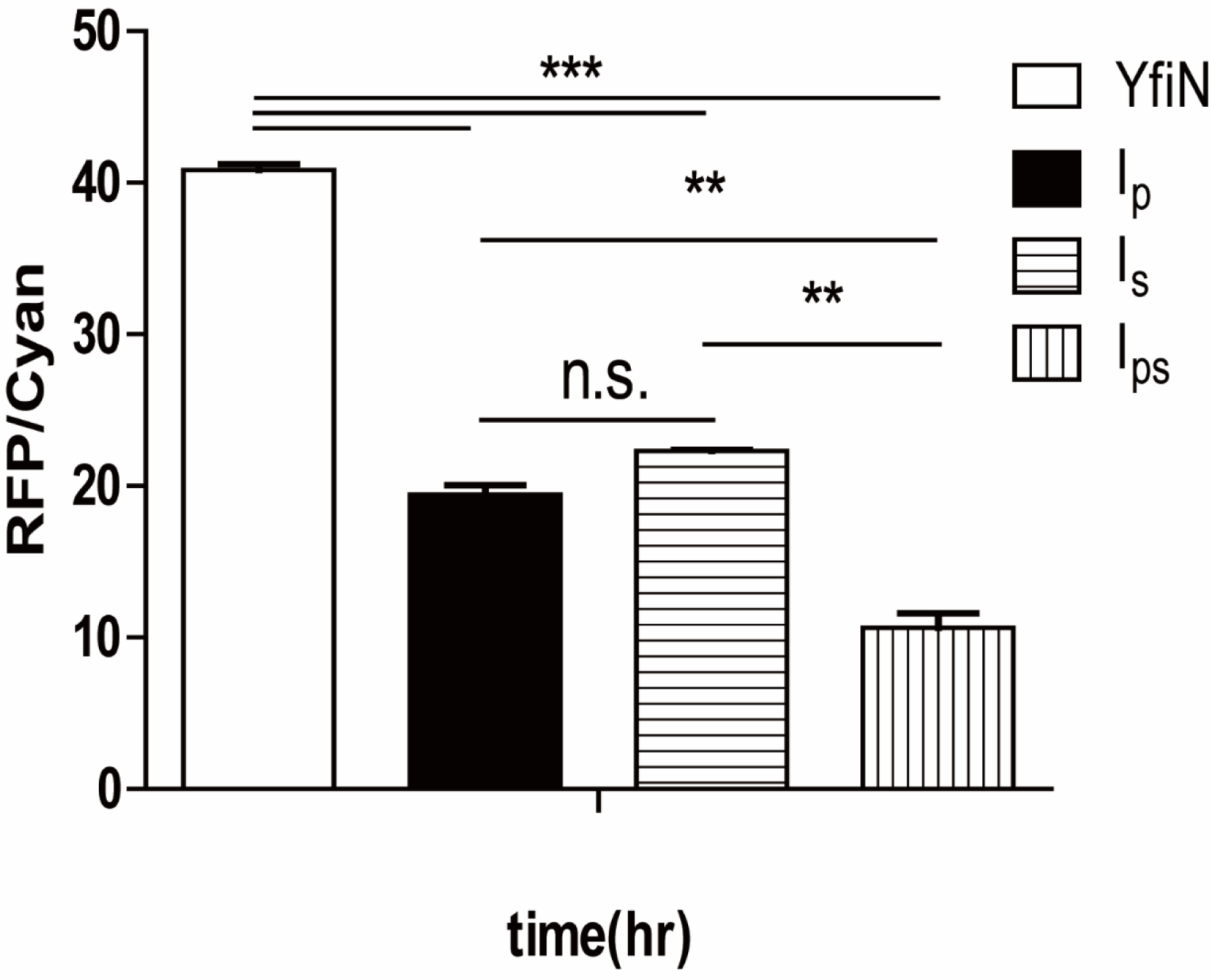
Comparison of c-di-GMP levels in cells expressing pYfiN and its I site mutants Ip, Is, Ips. The data were obtained by using a riboswitch-based fluorescent biosensor as described in Fig. S3B.

**Figure S5.**
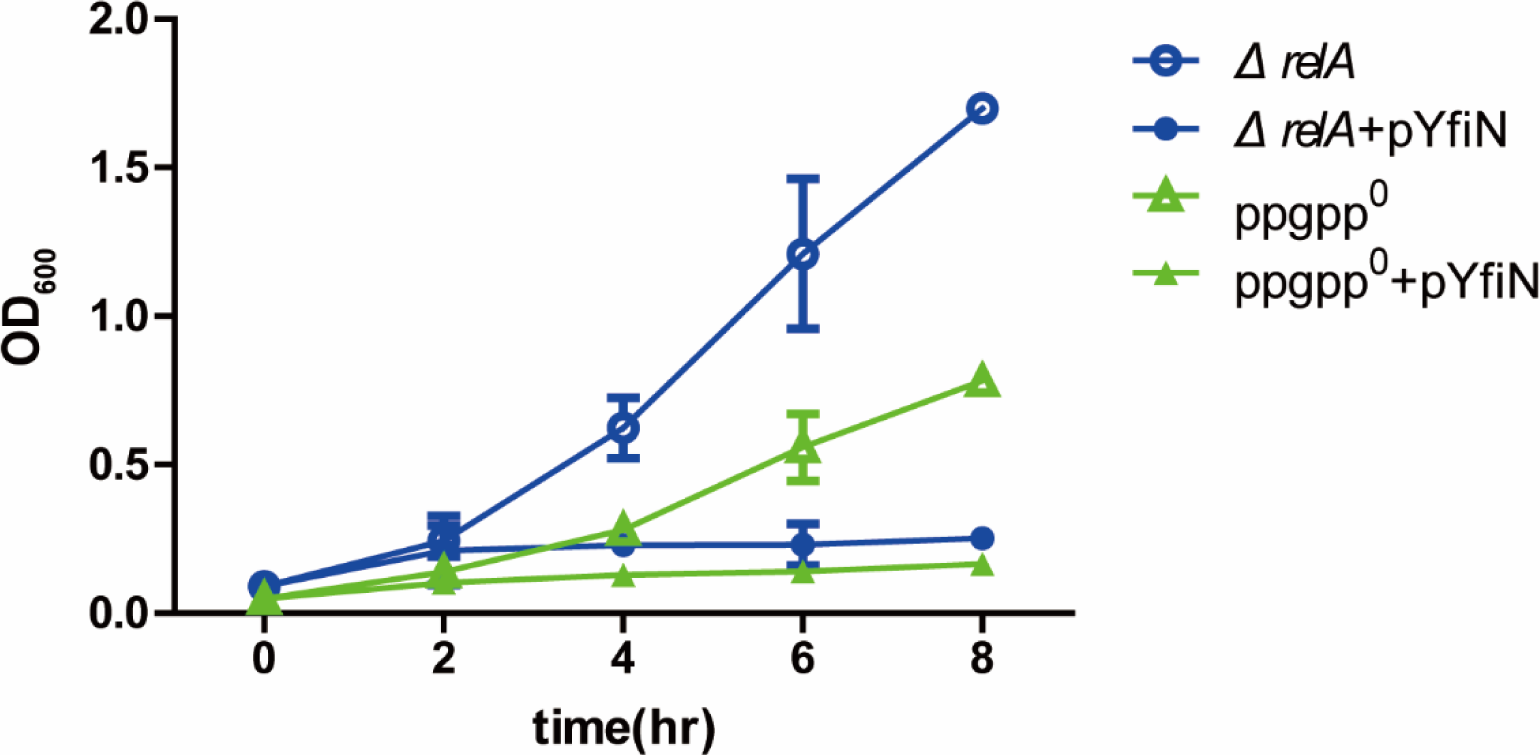
The (p)ppGpp pathway does not contribute to YfiN-mediated growth arrest. Time course growth of *ΔrelA* or *ΔrelAΔspoT*(ppGpp^0^) strains transformed with pYfiN and propagated in M9M with 0.02% arabinose.

**Figure S6.**
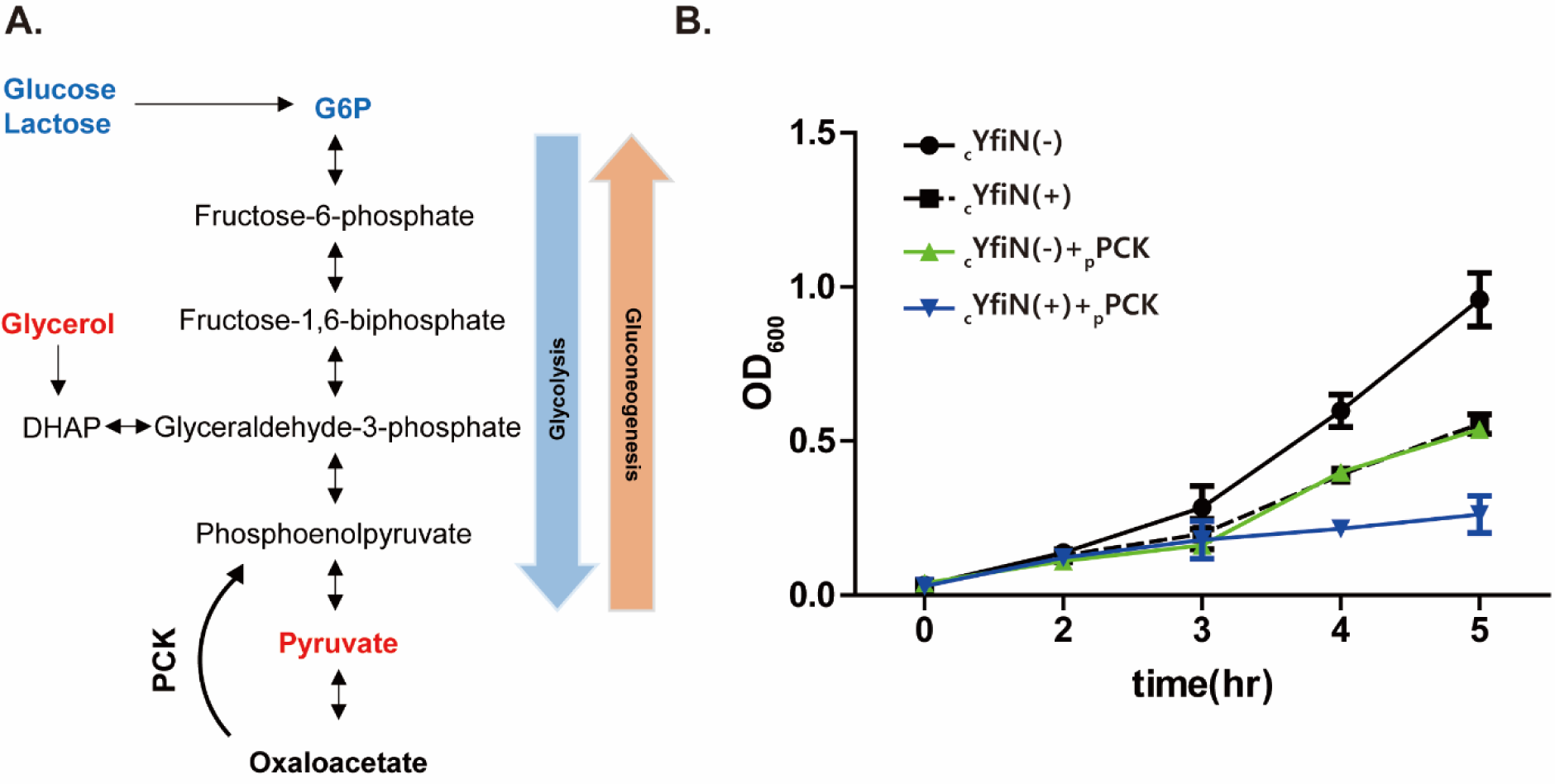
Co-expression of PCK with cYfiNGFP increases severity of growth arrest. *(A*) Gluconeogenesis pathway indicating the step at which PCK acts. *(B)* Growth curve in M9 mannitol of the cYfiN**GFP** (induced with arabinose: -/+) and strain transformed with pPCK (induced with IPTG).

**Figure S7.**
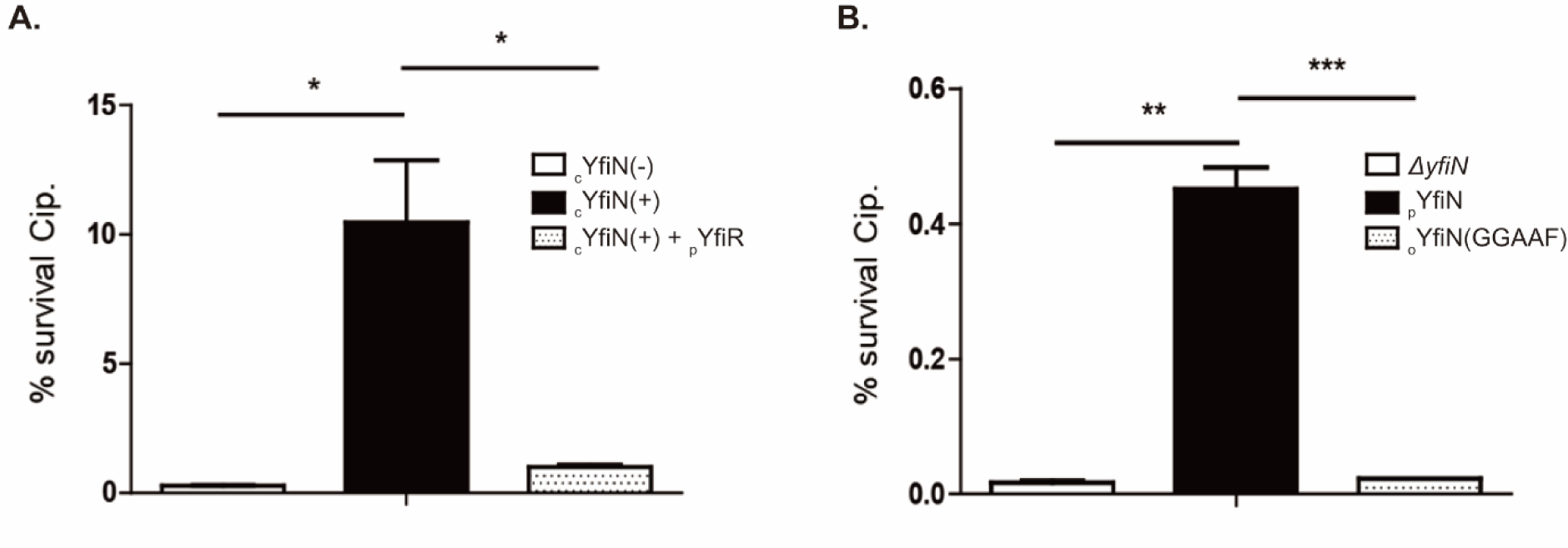
The DGC activity of YfiN is essential for antibiotic tolerance. (*A*) The cYfiNGFP strain was transformed with pYfiR (periplasmic inhibitor of YfiN) and induced with IPTG or arabinose, respectively. Cells were propagated in M9M for 4 hours before addition of Cip (20μg/ml) to the media. After incubation for 4 hours, % survival was determined by measuring CFU counts before and after antibiotic exposure. (*B*) As in *A* except with pYfiN and its active site mutants. The lower cell survival in B vs A is likely due to higher GTP depletion in pYfiN compared to cYfiN, leading to higher toxicity and cell death [58].

**Figure S8.**
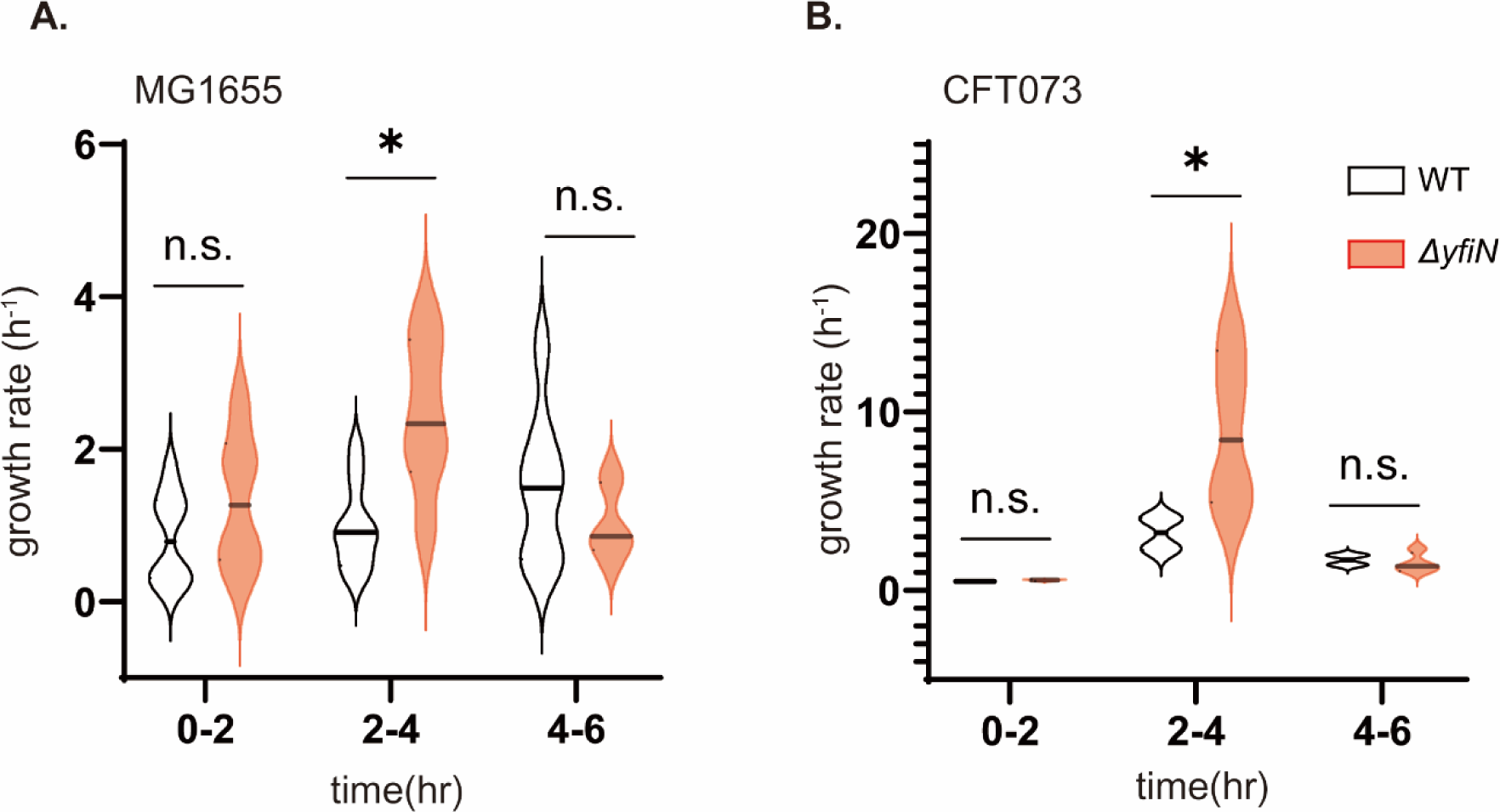
Growth rate comparisons of two *E. coli* WT strains and their *ΔyfiN* derivatives in M9 glycerol. *(A)* Data for the WT lab strain MG1655 are taken from Fig. 6A (n=8). *(B*) Similar experiments, but with the uropathogenic strain CFT073 (n=4).

**Figure S9.**
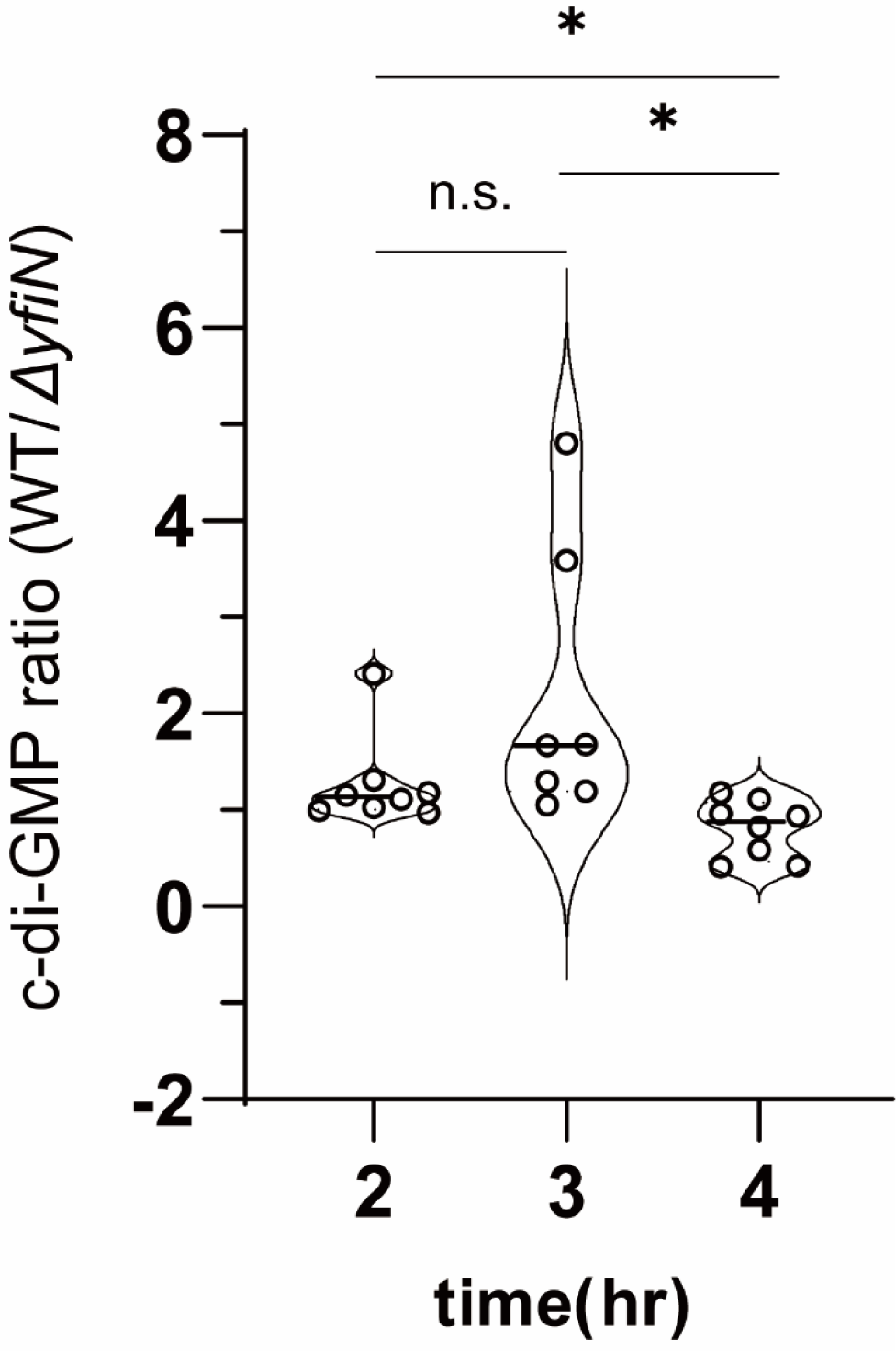
c-di-GMP levels in WT vs. ΔYfiN measured by the riboswitch-based biosensor. The strains were propagated in M9M and fluorescence measured as in Fig. S3B. Samples collected at indicated time points were diluted to OD600 of 0.5, prior to measurement. Each dot represents one biological replicate (n=8 for 2 & 4 hours, and 7 for 3 hours).

**Figure S10.**
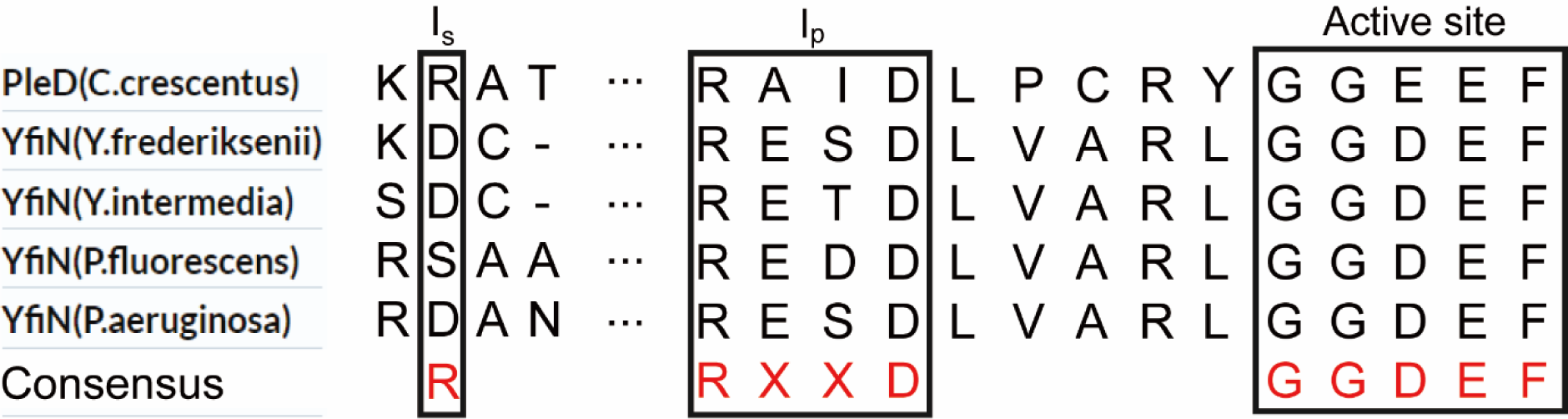
YfiN homologs that have Ip but not Is sites. The Ip site (RXXD), the predicted Is site (R), and the DGC active site (GGDEF) are boxed in a sequence alignment of YfiN from *Yersinia* (Y) and *Pseudomonas* (P) species, and compared to PleD from *C. crescentus*, which has a functional set of both I sites (see Fig. 3B).

**Table S1.**

| Strains | Genotype or Description | Reference |
| --- | --- | --- |
| MG1655 | K12 wild-type (WT) strain: F<sup>-</sup>, λ<sup>-</sup>, rph-1 | Laboratory Collection |
| HK359 | MG1655 $\Delta yfiN$ | [24] |
| HK461 | HK359 + pBAD30_YfiN<sub>GFP</sub> | [24] |
| JH101 | HK359 + pBAD30_YfiN | This study |
| HK532 | MG1655 yfiR::kan(←) yfiN::pBAD-yfiN-gfp | [24] |
| HK533 | MG1655 yfiR::kan(←) yfiN::pTrc-yfiN-gfp | This study |
| HK800 | HK359 + pBAD33_YfiN<sub>GFP</sub> |  |
| JH102 | MG1655 + pBAD33_DgcA | This study |
| JH100 | HK359 + pBAD30_YfiN | This study |
| JH103 | HK359 + pBAD30_YfiN(GGAFF) | This study |
| JH104 | HK359 + pBAD30_YfiN(I<sub>p</sub>) | This study |
| JH105 | HK359 + pBAD30_YfiN(I<sub>s</sub>) | This study |
| JH106 | HK359 + pBAD30_YfiN(I<sub>ps</sub>) | This study |
| JH107 | MG1655 + pBAD33_DgcA(GESD) | This study |
| JH108 | HK359 + pBAD33_YfiN(R260A)<sub>GFP</sub> | This study |
| JH109 | HK532 + ASKA_PCK | This study |
| JH202 | MG1655 + pBAD33_DgcC | This study |
| JH203 | MG1655 + pBAD33_DgcE | This study |
| JH204 | MG1655 + pBAD33_DgcF | This study |
| JH201 | MG1655 + pBAD33_DgcJ | This study |
| JP1932 | AW405 $\Delta bcsA$ csgD pgaC fimA wcaD yjbE | [32] |
| JH110 | HK359 + pBAD33_YfiN + pTrc99a_YhjH | This study |
| JH111 | HK532 + pTrc99a_YfiR | This study |
| JH250 | HK359 + pBAD30_YfiN-Flag (c-term.) | This study |
| JH251 | HK359 + pBAD30_YfiN(I<sub>ps</sub>)-Flag (c-term.) | This study |
| JH252 | HK359 + pBAD30_YfiN(GGAFF)-Flag (c-term.) | This study |
| JH253 | HK359 + pBAD30_YfiN(R260A)-Flag (c-term.) | This study |
| JH300 | HK359 + pBAD30_YfiN-Flag(c-term.) | This study |
| JH301 | HK359 + pBAD30_YfiN + pFY4535 | This study |
| JH302 | HK359 + pBAD30_YfiN(I<sub>p</sub>) + pFY4535 | This study |
| JH303 | HK359 + pBAD30_YfiN(I<sub>s</sub>) + pFY4535 | This study |
| JH304 | HK359 + pBAD30_YfiN(I<sub>ps</sub>) + pFY4535 | This study |
| JH305 | MG1655 + pFY4535 | This study |
| JH306 | HK359 + pFY4535 | This study |
| JH307 | JH110 + pFY4535 | This study |
| JH308 | HK359 + pBAD33_YfiN + pTrc99a_Empty + pFY4535 | This study |
| JH309 | JH102 + pFY4535 | This study |
| CFT073 | Uropathogenic strain of *E. coli* | R. Welch, U. Wisconsin-Madison |
| HK701 | CFT073Δ*yfiN* | Hyo Kyung Kim |
| Plasmids | Expressed Proteins | Reference |
| pKD4 | Kanamycin resistance gene template | [66] |
| pKD46 | λ Red Recombinase | [66] |
| pCP20 | FLP recombinase | [66, 69] |
| pTrc99a | Cloning vector, P<sub>Trc</sub>; Amp<sup>R</sup> | [69, 70] |
| pBAD30 | Cloning vector; P<sub>BAD</sub>; Amp<sup>R</sup> | [70] |
| pBAD33 | Cloning vector; P<sub>BAD</sub>; Cm<sup>R</sup> | [24, 70] |
| pBAD30_YfiN<sub>GFP</sub> | pBAD::YfiN-GFP(C-terminal) | [24] |
| pBAD30_YfiN | pBAD::YfiN | This study |
| pBAD33_YfiN | pBAD::YfiN | This study |
| pBAD30_YfiN(GGAAF) | pBAD::YfiN (D329A E330A) | This study |
| pBAD30_YfiN(I<sub>p</sub>) | pBAD::YfiN (G317R H320D) | This study |
| pBAD30_YfiN(I<sub>s</sub>) | pBAD::YfiN (N280R) | This study |
| pBAD30_YfiN(I<sub>ps</sub>) | pBAD::YfiN (N280R G317R H320D) | This study |
| pBAD33_DgcA | pBAD::DgcA | This study |
| pBAD33_DgcA(GESD) | pBAD::DgcA (R80G) | This study |
| pBAD30_YfiN(R260A)<sub>GFP</sub> | pBAD::YfiN (R260A)-GFP | Hyo Kyung<br>Kim |
| pBAD30_YfiN-Flag (c-term.) | pBAD::YfiN-Flag | This study |
| pBAD30_YfiN(I<sub>ps</sub>)-Flag (c-term.) | pBAD:: YfiN(I<sub>ps</sub>)-Flag | This study |
| pBAD30_YfiN(GGAFF)-Flag (c-term.) | pBAD:: YfiN(GGAFF)-Flag | This study |
| pBAD30_YfiN(R260A)-Flag (c-term.) | pBAD:: YfiN(R260A)-Flag | This study |
| pFY4535_pmmb-bc3-5-phok-sok | pMMBGent::c-di-GMP sensor | [34] |
| pBAD33_DgcC | pBAD::DgcC | This study |
| pBAD33_DgcE | pBAD::DgcE | This study |
| pBAD33_DgcF | pBAD::DgcF | This study |
| pBAD33_DgcJ | pBAD::DgcJ | This study |
| pTrc99a_Empty | pTrc99a::Empty | This study |
| pTrc99a_YhjH | pTrc::YhjH | This study |
| pTrc99a_YfiR | pTrc::YfiR | This study |

